# Cryo-EM Structures and Regulation of Arabinofuranosyltransferase AftD from Mycobacteria

**DOI:** 10.1101/2019.12.22.885152

**Authors:** Yong Zi Tan, Lei Zhang, José Rodrigues, Ruixiang Blake Zheng, Sabrina I. Giacometti, Ana L. Rosário, Brian Kloss, Venkata P. Dandey, Hui Wei, Richard Brunton, Ashleigh M. Raczkowski, Diogo Athayde, Maria João Catalão, Madalena Pimentel, Oliver B. Clarke, Todd L. Lowary, Margarida Archer, Michael Niederweis, Clinton S. Potter, Bridget Carragher, Filippo Mancia

## Abstract

*Mycobacterium tuberculosis* causes tuberculosis, a disease that kills over one million people each year. Its cell envelope is a common antibiotic target and has a unique structure due, in part, to two lipidated polysaccharides – arabinogalactan and lipoarabinomannan. Arabinofuranosyltransferase D (AftD) is an essential enzyme involved in assembling these glycolipids. We present the 2.9 Å resolution structure of *M. abscessus* AftD determined by single particle cryo-electron microscopy. AftD has a conserved GT-C glycosyltransferase fold and three carbohydrate binding modules. Glycan array analysis shows that AftD binds complex arabinose glycans. Additionally, AftD is non-covalently complexed with an acyl carrier protein (ACP). 3.4 and 3.5 Å structures of a mutant with impaired ACP binding reveal a conformational change that suggests the ACP may regulate AftD function. Using a conditional knock-out constructed in *M. smegmatis*, mutagenesis experiments confirm the essentiality of the putative active site and the ACP binding for AftD function.

## INTRODUCTION

Globally, tuberculosis (TB) is the leading cause of death from a single infectious agent, out-ranking even HIV/AIDS (WHO, 2018). The causative agent of TB is *Mycobacterium tuberculosis*, which belongs to a genus with a unique cell envelope made of lipids and carbohydrates (Figure 1A). The major component of the mycobacterial cell envelope is a large macromolecule consisting of covalently linked peptidoglycan (PG), branched heteropolysaccharide arabinogalactan (AG) and long chain (70 to 90 carbon atoms long) mycolic acids, termed the mycolyl-arabinogalactan-peptidoglycan (mAGP) complex (Alderwick et al., 2015; Grzegorzewicz et al., 2016). This mAGP complex is in turn decorated by a variety of glycolipids called phosphatidyl-*myo*-inositol mannoside (PIM), lipomannan (LM) and lipoarabinomannan (LAM) (Figure 1A). *M. tuberculosis* produces between 2,000 to 10,000 distinct lipids (Layre et al., 2014), most of which form an outer membrane of very low permeability (Hoffmann et al., 2008; Nikaido and Jarlier, 1991). This cell envelope is crucial for growth and virulence of *M. tuberculosis* (Barry III, 2001; Jankute et al., 2015) and is a major contributor to the resistance of *M. tuberculosis* to common antibiotics. The importance of this unique structure in mycobacteria makes its biosynthetic enzymes attractive drug targets (Abrahams and Besra, 2016; Barry et al., 2007). For example, isoniazid and ethambutol, two of the four TB front-line drugs, target mycobacterial cell wall synthesis (Islam et al., 2017).

**Figure 1.**
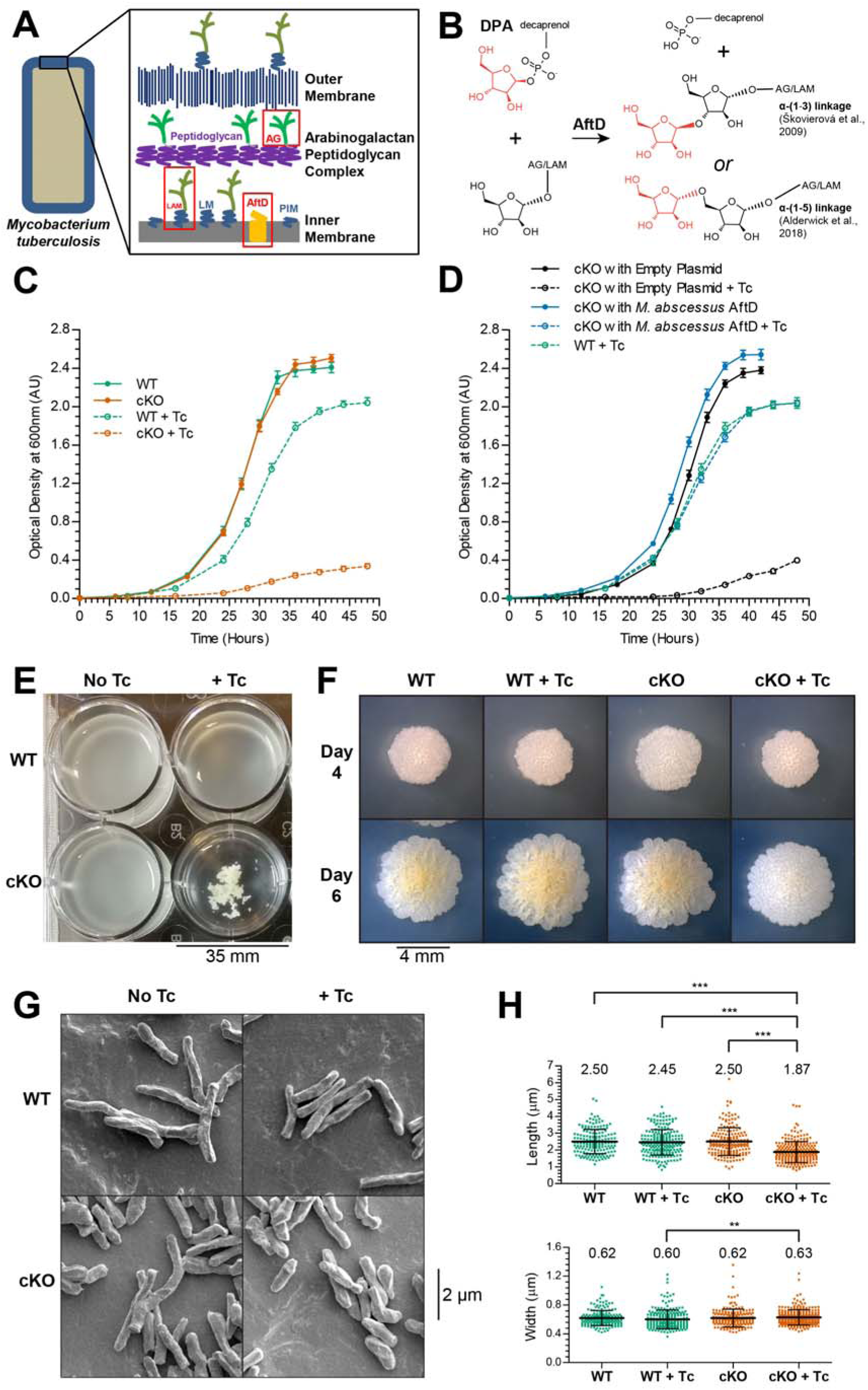
Functional analysis of the role of the arabinofuranosyltransferase AftD in the biosynthesis of the mycobacterial cell envelope. (A) Model of the cell envelope of *M. tuberculosis*. Red boxes highlight the arabinogalactan (AG) and lipoarabinomannan (LAM) components which are synthesized by AftD (yellow). (B) Reaction catalyzed by AftD enzyme. (C) Growth of wild-type *M smegmatis* (WT) and the conditional *aftD* mutant ML2218 (cKO) without and with 20 ng/mL tetracycline. (D) Complementation of the cKO with empty plasmid and plasmid carrying the *M. abscessus aftD* gene. The strains were grown with and without tetracycline. Both growth curves were recorded in biological triplicates. Standard deviations are plotted. (E) Clumping of the cKO cells in the presence of 20 ng/mL tetracycline in liquid culture in microplates. (F) Colony morphology of WT and cKO *M smegmatis* without and with 20 ng/mL tetracycline grown at 37 °C for 4 and 6 days. (G) Scanning electron microscopy micrographs and quantification of the cell length and width (H) from these images. For the cell lengths and widths (H), one-way ANOVA Kruskal-Wallis parametric test was done as the distribution was not normal. Dunns post-test was done between all pairs of samples.

Of the enzymes involved in mAGP biosynthesis, arabinofuranosyltransferases are responsible for the addition of D-arabinofuranose sugar moieties to AG and LAM (Abrahams and Besra, 2016). All mycobacterial arabinofuranosyltransferases are predicted to be multi-pass transmembrane (TM) proteins that use decaprenylphosphoryl-D-arabinofuranose (DPA) to donate an arabinofuranose residue to the mAGP (Wolucka et al., 1994). Arabinofuranosyltransferases add 90 arabinofuranose residues to the galactan core of AG and 55–72 arabinofuranose residues to the mannan core of LAM. The best characterized arabinofuranosyltransferases are EmbA, EmbB and EmbC (Belanger et al., 1996; Telenti et al., 1997), which are named Emb because they are putative targets of ethambutol (Goude et al., 2009; Takayama and Kilburn, 1989). Other family members include AftA (Alderwick et al., 2006), AftB (Seidel et al., 2007), AftC (Birch et al., 2008), AftD (Škovierová et al., 2009), and possibly other yet to be identified enzymes (Angala et al., 2016). AG synthesis starts when AftA adds a single D-arabinofuranose to the C5 of the D-galactofuranose at positions 8, 10 and 12 of the existing galactan. Thereafter, EmbA and EmbB extend the arabinan chain through α-(1→5) linkages; α-(1→3) branching is then introduced by AftC and possibly AftD. AftB is the final enzyme to cap off the AG with addition of a terminal β-(1→2)-linked D-arabinofuranose. For LAM, an unknown enzyme (Angala et al., 2016) primes the mannan chain with an α-(1→5) linkages by EmbC. AftC and possibly AftD then catalyzes α-(1→3) branching and capping in a β-(1→2) manner by AftB, as in the synthesis of AG. These pathways are by no means fully characterized (Alderwick et al., 2006).

Arabinofuranosyltransferase D (AftD) is essential for growth (Škovierová et al., 2009) and, with a theoretical molecular mass of 150 kDa, it is the largest predicted glycosyltransferase encoded in the mycobacterial genome. It is described as being involved in either α-(1→3) (Škovierová et al., 2009) and/or α-(1→5) (Alderwick et al., 2018) branching activity, and is thought to add the final few α linked arabinofuranose residues to AG and LAM (Alderwick et al., 2018) (Figure 1B). Little is known about both the function and structure of AftD, although its large size in comparison to the other arabinofuranosyltransferases (which range from 49 to 118 kDa) has raised the question of whether it has additional functions beyond catalyzing glycosidic bond formation (Škovierová et al., 2009).

To better understand the function of this enzyme, we determined the 2.9 Å structure of recombinant AftD from *M. abscessus* produced in *E. coli* using single-particle cryogenic electron microscopy (cryo-EM). We analyzed its *in vitro* carbohydrate binding capability via glycan array analysis and we constructed a conditional *aftD* deletion mutant in *M. smegmatis* to interrogate the functionality of AftD mutants *in vivo*. Unexpectedly, analysis of the structure revealed that AftD is tightly bound to an acyl carrier protein (ACP). The physiological relevance of this interaction was confirmed in *smegmatis* through mass spectrometry. Finally, we purified and determined structures of an AftD mutant designed to abrogate ACP binding to 3.4 and 3.5 Å resolution. These structures revealed ordering of a loop at the putative active site, suggesting that the ACP may regulate AftD function.

## RESULTS

### Phylogenetic Analysis Reveal Existence of AftD Homologs Across *Actinobacteria*

Iterative rounds of PSI-BLAST (Altschul and Koonin, 1998) were used to identify homologs of AftD using the amino acid sequence of the protein from *M. tuberculosis* as a template. This analysis revealed that AftD is found not just in the *Corynebacteriales* order, but also in 11 other orders of the *Actinobacteria* class (Ludwig et al., 2015) including *Streptomycetales*, *Propionibacteriales* and *Frankiales* (Figure S1). The homologs all have a similar predicted topology, computed using the TMHMM server (Krogh et al., 2001), with an N-terminal TM region, followed by a long periplasmic region, and a shorter C-terminal TM region. The average length of all AftD proteins is 1381 amino acids. Most sequences (including the mycobacterial ones) cluster around this length, while AftD homologs in corynebacteria are significantly shorter with an average length of 1070 residues (Figure S2A). Excluding corynebacteria, the average length of AftD from the remaining genera is 1422 residues.

### Conditional Deletion of the *M. smegmatis aftD* Gene Results in Altered Cell Phenotype

As AftD has been reported to be an essential enzyme in *M. tuberculosis* (DeJesus et al., 2017) and in *M. smegmatis* (Škovierová et al., 2009), a conditional knock-out was designed to interrogate the function of AftD in *M. smegmatis* and provide a system for structure-based mutagenesis studies. First, we constructed the *M. smegmatis* ML2217 strain (WT), which constitutively produces the reverse Tet repressor TetR38. Then, the chromosomal *aftD* gene in *M. smegmatis* ML2217 was placed under the control of the tetO-4C5G operator (Kim et al., 2013), which is constitutively transcribed in the absence of tetracycline (Figure S2B). In the presence of tetracycline, the TetR38 binds to tetO-4C5G, repressing transcription of the *aftD* gene. To our knowledge this is the first conditional AftD knock-out in mycobacteria that can be readily controlled in a temperature-independent fashion.

Phenotypic analyses of wild-type *M. smegmatis* (WT, ML2217) and the conditional *aftD* knock-out mutant (cKO, ML2218) were performed in medium with and without tetracycline (Figure 1). When grown in 7H9 medium without tetracycline, both WT and cKO strains had similar growth curves, reaching an OD_600nm_ of 2.5 after about 32 hours (Figure 1C). When 20 ng/mL of tetracycline was added to the growth media, OD_600nm_ failed to reach 0.5 after 48 hours (Figure 1C), confirming that *aftD* is indeed essential for growth of *M. smegmatis* under those conditions. In the presence of tetracycline to silence *aftD* expression, cKO cells readily aggregated under low shaking speed, suggesting the properties of the cell surface have changed (Figure 1E). The colony morphology of the conditional *aftD* mutant was also altered, appearing less rough than the controls (Figure 1F). When imaged by scanning electron microscopy, cells from the cKO strain appeared shorter in the presence of tetracycline (Figure 1G). Indeed, this observation was corroborated by measurements of the cell length and width, showing a statistically significant reduction in cell length of the cKO strain in the presence (1.9 ± 0.6 μ) versus the absence (2.5 ± 0.8 μ) of tetracycline, while the widths remained the same (Figure 1H).

### Genomic Expansion and Complementation of the *M. abscessus aftD* Gene

To study the AftD function *in vitro* and determine its structure, we adopted a structural genomics approach (Bruni and Kloss, 2013; Love et al., 2010; Mancia and Love, 2010) to identify homologs that yielded high expression levels in *E. coli* and were stable in detergents compatible with structure determination (see Methods). High-throughput small-scale expression and purification screens of AftD from 14 mycobacterial species revealed that under our experimental conditions only the *M. abscessus* homolog was expressed and purified from *E. coli* in quantities detectable in Coomassie-stained protein gels. This homolog has a high sequence identity of 62% and 70% with the *M. tuberculosis* and *M. smegmatis* AftDs, respectively.

To confirm the *in vivo* functionality of *M. abscessus* AftD, we complemented the cKO with the *M. abscessus aftD* gene. Initial experiments using tetracycline or acetamidase promoters to express the *aftD* gene resulted in cell aggregation, suggesting altered cell surface properties of *M. smegmatis*, probably resulting from *aftD* overexpression. However, an expression plasmid containing the *M. abscessus aftD* gene transcribed from its endogenous promoter (located in a 200 base pair upstream fragment) fully restored wild-type growth of the cKO strain in the presence of tetracycline (Figure 1D). From these results we could conclude that *M. abscessus* AftD is functional in *M. smegmatis*, and that the phenotype observed in the conditional *M. smegmatis* mutant is indeed due to the depletion of the *M. smegmatis* AftD protein.

### Structure Determination Using Cryo-EM

We determined the structure of the full-length *M. abscessus* apo-state AftD using single-particle cryo-EM. Despite substantial efforts to optimize the over-expression and purification of AftD in *E. coli*, final yields after purification were low: less than 100 μ of protein could be isolated from 9.6 L of cell culture. AftD was successfully incorporated into lipid-filled nanodiscs after a two-step purification in detergent (Figures S3A and S3B). To generate multiple cryo-EM grids of the sample at various conditions for screening and data collection despite the low yield, a picoliter dispensing robot (Spotiton) was used to vitrify the sample (Dandey et al., 2018; Jain et al., 2012; Razinkov et al., 2016). Data were collected on a Titan Krios microscope equipped with a Gatan K2 Summit camera fitted with an energy filter. 7,274 micrographs were collected and 889,985 particles were initially processed (Figure S3C and Table S1). After data processing, the final map was produced from 490,616 particles, resulting in a reconstruction with an overall resolution of 2.9 Å (Figure S3). An atomic model for the entire protein was built, with the exception of both termini (residues 1 and 1392-1410) and some disordered loop regions (253-259, 270-286, 330-356, 390-401, 1281-1285, 1358-1367) (Figure 2). The region represented by residues 950-1090 was flexible; signal subtraction followed by focused classification of this part (Bai et al., 2015) (see Methods) improved the map quality to allow for better model building. The final refined model exhibits good stereochemistry and correlates well with the density map (Figure S4 and Table S1).

**Figure 2.**
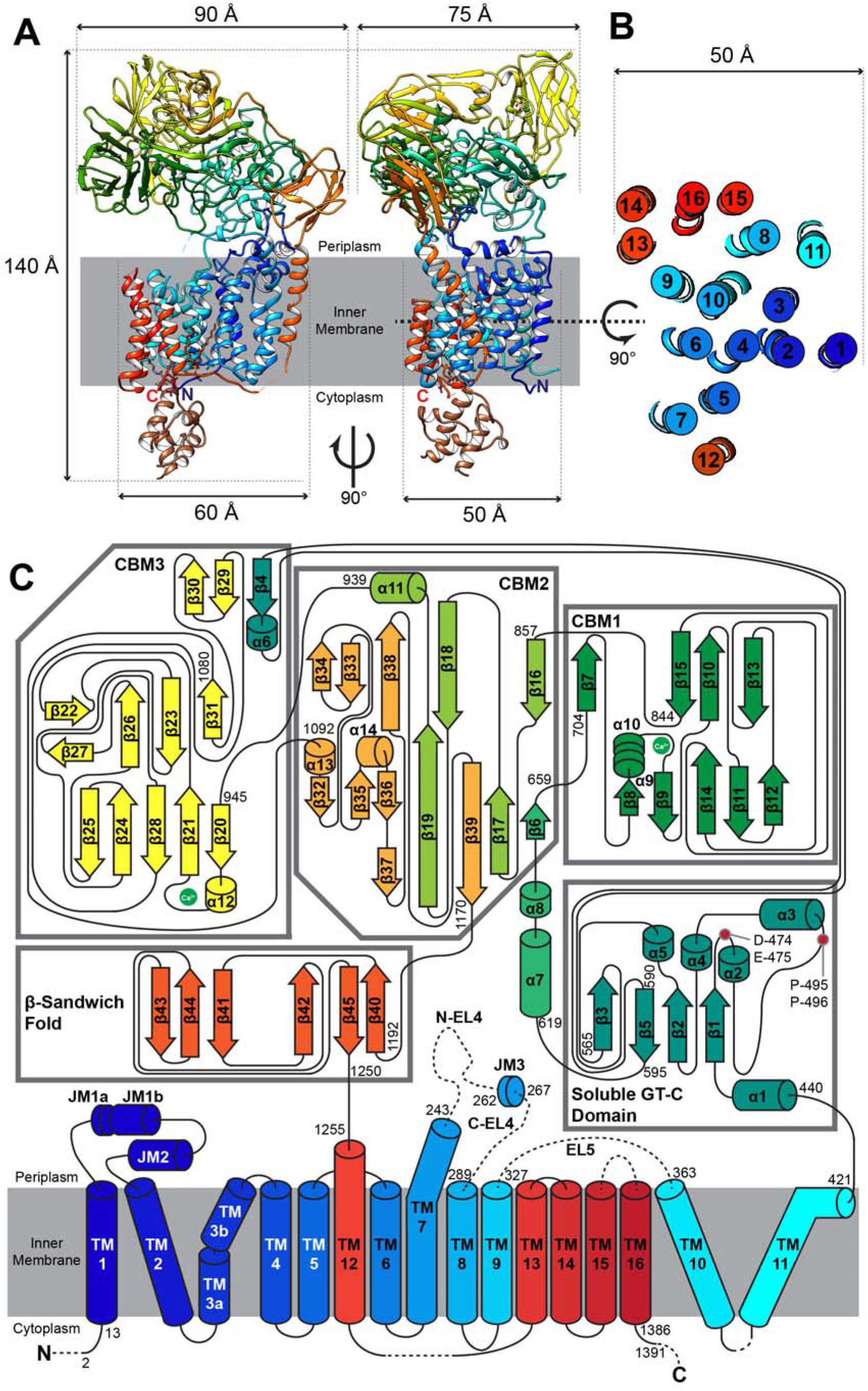
Architecture of AftD Complex. (A) Single-particle cryo-EM structure of AftD complex, rendered in cartoon and colored in rainbow from N terminus (blue) to C terminus (red). The complexed *E. coli* acyl carrier protein and ligands are colored in brown. Both proteins are rendered as cartoon. Ca^2+^ ions are rendered as green spheres. Two orthogonal views that are perpendicular to the plane of the membrane are shown. The approximate dimensions of the monomer are 85, 145, and 75 Å (width, height, and depth). Membrane boundaries were derived from the interface between the nanodisc lipids and solvent. (B) Transmembrane helices arrangement of AftD, viewed as a slice and magnified. (C) Two-dimensional topological diagram of AftD. The individual soluble domains are enclosed in separate grey boxes. The topology diagram is colored in rainbow from N terminus (blue) to C terminus (red). Unbuilt parts of the model due to poor map density are indicated by dotted lines. The two bound Ca^2+^ atoms are shown as green circles, and side chains coordinating them are shown as sticks. The putative catalytic residues previously hypothesized (Škovierová et al., 2009) are shown as red circles.

### Architecture of AftD

AftD consists of a membrane embedded portion with 16 TM helices and a soluble portion that resides on the periplasmic side (Hoffmann et al., 2008) of the inner membrane (Figure 2A). Both termini of AftD project into the cytoplasm. Starting from the N-terminus, the first 11 TM helices form a typical GT-C glycosyltransferase fold (Figures 2A and 2B) (Liu and Mushegian, 2003). Thereafter, the polypeptide chain enters the periplasm. The soluble, periplasmic portion of AftD is large, accounting for ∼60% of the mass of the protein, and is made up of five distinct domains: a domain of mixed α β folds that is homologous to other solved GT-C structures (discussed below), three carbohydrate binding modules (CBMs) and one structural β-sandwich domain (Figure 2C). Though these domains are discrete, they have extensive inter-domain interactions that maintain them tightly associated, including some secondary structural elements: β4 from the soluble GT-C domain forms a beta-sheet with β29 and β30 of CBM3, β7 of CBM1 forms a beta-sheet with β CBM2 (Figure 2C). After the soluble domains, the polypeptide chain enters back into the membrane via a single TM helix (TM12) that connects via a long loop to TMs 13-16 and the C-terminus as a separate membrane-bound domain from the N-terminal TM helices (Figure 2C).

### AftD Has a Conserved Glycosyltransferase GT-C Fold

Using the Dali server (Holm and Laakso, 2016), atomic structures with structural similarity to AftD were retrieved and top hits corresponded to all full-length GT-C structures deposited in the database: yeast oligosaccharyltransferase STT3 (PDB ID: 6EZN, 6C26) (Bai et al., 2018; Wild et al., 2018), *Archaeoglobus fulgidus* oligosaccharyltransferase AglB (PDB ID: 3WAJ, 5GMY, 3WAK) (Matsumoto et al., 2013; Matsumoto et al., 2017), *Campylobacter lari* oligosaccharyltransferase PglB (PDB ID: 3RCE, 5OGL, 6GXC) (Lizak et al., 2011; Napiórkowska et al., 2018; Napiórkowska et al., 2017), *Cupriavidus metallidurans* aminoarabinose-transferase ArnT (PDB ID: 5EZM, 5F15) (Petrou et al., 2016) and recently yeast mannosyltransferase Pmt1-Pmt2 (PDB ID: 6P2R, 6P25) (Bai et al., 2019). The first ∼600 residues of AftD correspond to this homologous GT-C fold, and will subsequently be referred to as the GT-C super-domain of AftD (Figures 2C and S5C). This GT-C super-domain comprises a total of 11 TM helices (colored blue in Figure 2B) – two less than all the other solved GT-Cs (STT3, AglB, PglB and ArnT). The two missing TM helices are also not conserved amongst GT-Cs (Petrou et al., 2016) and correspond to TM helix 8 for STT3 (Figure S5D), helix 8 and 9 for PglB (Figure S5E), and helix 12 and 13 for ArnT (Figure S5F). The remaining 11 TM helices are structurally homologous to the other solved GT-Cs in several aspects: 1) They are in the same arrangement in AftD as seen in the other four solved GT-C structures; 2) There are two periplasmic juxtamembrane (JM) helices (JM1 and JM2) between TM1 and TM2; 3) TM3 is kinked by a proline and broken into two smaller helices; 4) TM7 extends beyond the membrane into the periplasm and is joined by a longer loop to TM8. A short alpha helical JM helix (JM3) is present in the loop; 5) A soluble periplasmic domain is always present, composed of an α /α arrangement (Figure 2C).

### Putative Active Site and Substrate Binding Pocket

The postulated substrates for AftD are the donor DPA (Alderwick et al., 2007) and a poly-arabinose acceptor that is part of the growing AG or LAM (Škovierová et al., 2009) (Figure 1B). Although the structure of AftD was solved in the apo state, the high degree of homology as described above between AftD’s GT-C super-domain and the other solved GT-C glycosyltransferases allows for inference of mechanistic insights. In the GT-C super-domain, a large cavity is present between the periplasmic domain and the periplasm-facing ends of TM4, 6, 7, 9 and 10, with a volume of ∼1240 Å^3^ (Figure 3A). Using the unique AftD sequences of *Actinobacteria* class, sequence conservation was mapped onto the structure of AftD to reveal that this cavity contains a number of highly conserved residues, including E251, D474 and E475, which are the most plausible catalytic residue candidates. When representative eukaryotic (STT3) and bacterial (PglB) structures were aligned with the GT-C super-domain, the active sites of both structures are in the same area as that of AftD (Figure 3B). Other GT-C glycosyltransferases like PglB require the use of divalent metal cations to function (Sharma et al., 1981), and AftD seems to have a putative cation binding pocket (Figures 3A and 3B, red dotted line circle). Interestingly, the charge around the putative active site for AftD is slightly positive, compared to slightly negative for STT3 and PglB. This is consistent with the different acceptor substrates of these proteins: STT3 has preference for peptides with aromatic residues at the −2 position (Wild et al., 2018), PglB has preference for negatively charged side chain at the −2 position (Napiórkowska et al., 2017), while AftD binds to a poly-arabinofuranose glycan moiety (which can form ion-dipole interactions with positively charged residues).

**Figure 3.**
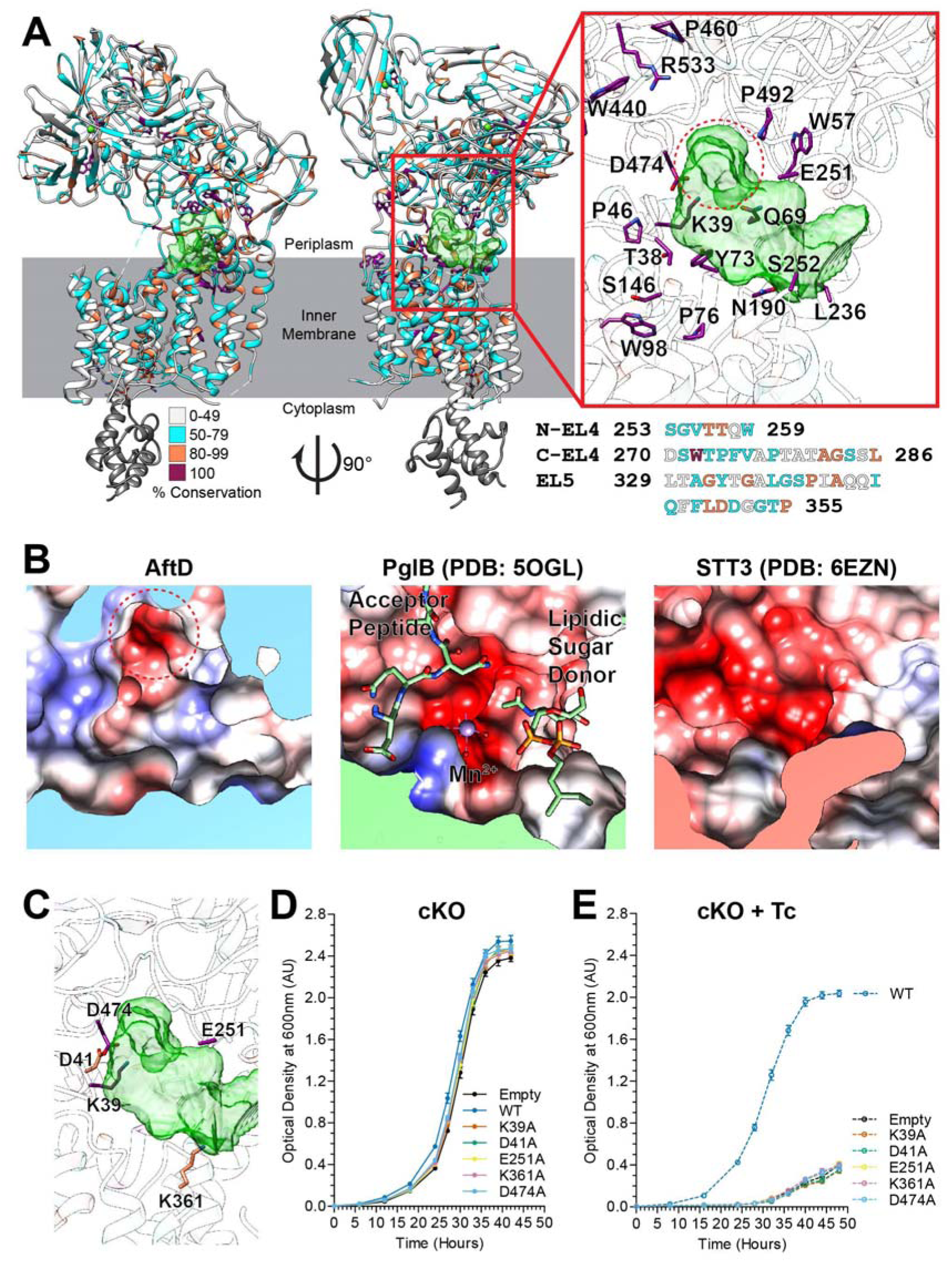
Sequence conservation and putative active site of AftD. (A) Structure of AftD, rendered in cartoon and colored based on sequence conservation. Unique sequences recovered from PSI-BLAST search were first pruned to those that are 95% the length of *M. abscessus* AftD (see Figure S3B). Thereafter, closely related sequences were removed by CD-HIT, giving a list of 289 sequences across the *Actinobacteria* class. The putative active site cavity, generated by Voss Volume Voxelator server (Voss and Gerstein, 2010) using probes of 2 and 5 Å radii, is colored in semi-transparent green. Residues with 100% conservation are colored in magenta and have their side chains displayed. The insert shows putative active site cavity with strictly conserved residues around it labeled. The primary sequences of the un-modeled flexible loops around the active site are shown below the insert, colored by conservation. (B) Columbic potential maps of the putative active site region of AftD, aligned PglB and yeast OST (STT3). The putative divalent cation binding pocket is circled with a red dotted line, and is mirrored in (A). The ligands and Mn^2+^ ion in the structure of PglB are shown as sticks and ball respectively. (C) Residues that have been mutated in the *M. abscessus aftD* gene have side chains displayed as stick, and the model colored by conservation. These mutant plasmids were then transformed into the cKO to assess the effect of these mutations on the growth rate of *M. smegmatis*, in the absence (D) or presence of tetracycline (E).

A shallow groove along TMs 6, 9 and 13 of AftD starts at the cytoplasmic side of the membrane and ends at the putative active site, and is likely to accommodate the putative DPA donor (Figure S5B). A diacyl-phospholipid (modeled as phosphatidylethanolamine) can be resolved in the density map near the cytoplasmic region. There is moderate conservation in this region, and it is also homologous to the polyprenol binding pockets of STT3 (Figure S5D), PglB (Figure S5E) and ArnT (Figure S5F), making this site the likely binding pocket for the lipidic section of DPA. In comparison with the locations of lipidic donor substrates in the other homologs, it seems that although the general binding area is conserved, the exact identities of the helices that form the binding site are likely to differ.

There are three long loops (N-EL4 between TM 7-JM 3, C-EL4 between JM 3-TM 8 and EL5 between TM 9-10) that do not have clear density present, and consist of 7, 17 and 27 residues, respectively (Figures 2C and 3A). Given that the structure of AftD was solved in the apo-state, these unresolved loops could be analogous to those of other glycosyltransferases that span across the active site and become ordered only upon ligand binding. Indeed, superimposition against the structures of STT3 (Figure S5D), PglB (Figure S5E) and ArnT (Figure S5F) suggests that the STT3 EL5, PglB EL5 and ArnT PL4 loops are analogous to the AftD EL4 loop, as they all extend from analogous TM helices. In contrast, AftD EL5 seems to be unique to this enzyme (Figures S5D, S5E and S5F). A number of conserved residues are present on both the loops, suggesting these segments are likely to be important for AftD’s enzymatic function. In particular, W272 is 100% conserved and D351 is moderately conserved (Figure 3A).

We designed a series of mutants based on the apo structure of AftD to examine structure-based functional hypotheses. Mutations around the putative active site were designed to target highly conserved residues that could be involved in substrate binding or catalysis (Figure 3C). These residues (K39, D41, E251, K361, D474) were mutated to alanine. All mutated AftD proteins were expressed in *E. coli* and purified using non-ionic detergents to yields comparable to WT, suggesting that the proteins produced were properly folded (Figure S6). All of these *aftD* mutations were also cloned in the mycobacterial expression vector to examine whether these mutated *aftD* genes restored WT growth of the conditional *M. smegmatis aftD* mutant with tetracycline. None of these AftD mutants were capable to complement the growth defect of the conditional *M. smegmatis aftD* mutant under non-permissive conditions. These results suggest that these residues play essential roles for proper functioning of AftD (Figures 3D and 3E).

### Carbohydrate Binding Modules of the AftD Soluble Domain Help Bind Complex Arabinofuranose Chains

In addition to the GT-C super-domain, AftD has a large soluble domain consisting of three carbohydrate binding modules (CBMs) that exhibit beta-sandwich folds (Figure 4A). There is an unusual structural motif of four consecutive prolines (residues 863-866) on CBM2, which appears to serve a structural role by making hydrophobic interactions with six other neighboring prolines from within CBM2 and also from a region between CBM1 and the GT-C soluble domain (Figure S5A). The structure shows that two CBMs (CBM1 and CBM3) bind an ion that is likely calcium, based on comparison with other structural homologs (PDB ID: 1UYY and 4A45). Notably, CBM3 is more flexible than the other two CBMs, as evidenced by the smearing of the cryo-EM density at CBM3 before focused classification.

**Figure 4.**
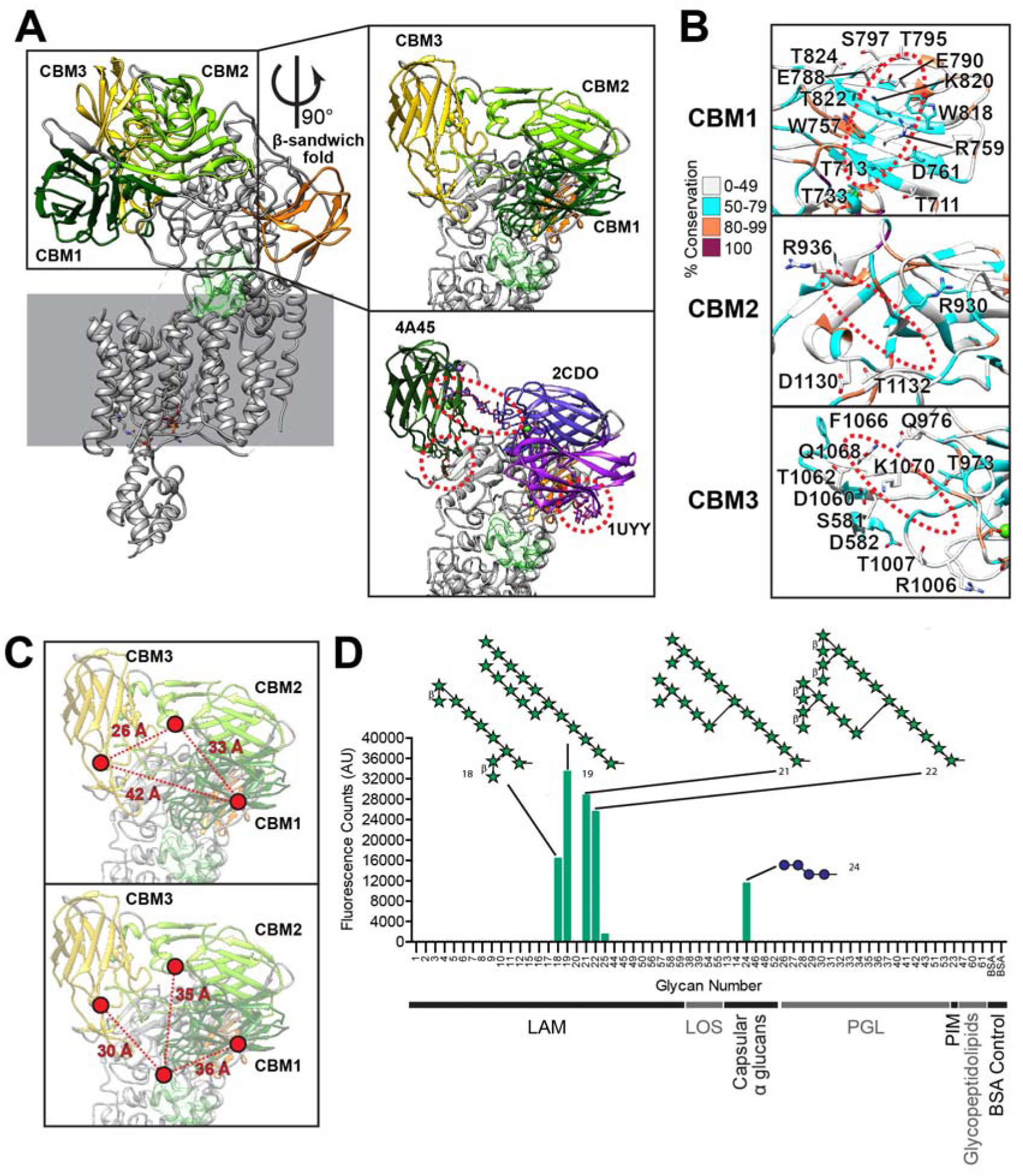
The Carbohydrate Binding Modules (CBMs) of AftD. (A) The three soluble CBMs and a β sandwich fold domain are colored in the AftD structure. The model is rendered as cartoon. In the first insert on top, the CBMs are zoomed in and the putative active site cavity is colored in semi-transparent green as in Figure 3. In the bottom insert, similar protein structures (PDB IDs indicated) aligned to and replacing each AftD CBM to show putative glycan binding scheme. (B) Putative sugar binding interfaces of the CBMs, with residues that could be involved in binding labelled and the putative location of the sugars circled in red dotted lines. All residues are colored by conservation. (C) Distances among the CBMs (top panel) and to the putative ligand binding cavity displayed in light green (bottom panel). (D) Glycan array analysis of AftD, with top glycan motif hits indicated, where green star represents arabinofuranose and blue sphere glucopyranose. The classes of glycans used are: lipoarabinomannan (LAM), trehalose mycolates and lipooligosaccharides (LOS), capsular α glucans, phenolic glycolipids (PGL), phosphatidyl*-myo-* inositol mannoside (PIM), glycopeptidolipid and bovine serum albumin (BSA) control. For structures of all glycans on the array see (Zheng et al., 2017a).

Structural homologs for each of the CBMs were identified using the Dali server and the highest scoring structural homolog with a ligand is shown in Figure 4A to illustrate putative glycan binding sites. The putative glycan binding sites of CBM1 and 3 have an extensive network of polar (S, T, N, Q) and charged (D, E, R, K) residues that can form ion-dipole or hydrogen bonds with the carbohydrate ligand (Figure 4B). In addition, CBM1 and CBM3 have aromatic side chains (W757, W818 for CBM1 and F1066 for CBM3) that can facilitate hydrophobic stacking interactions with the face of carbohydrate rings, an important mechanism of binding in CBMs (Boraston et al., 2004). These findings suggest that CBM1 and CBM3 are likely carbohydrate binding modules. CBM2, on the other hand, has only four polar/charged and no aromatic residues at the putative binding site indicating that it binds carbohydrates only weakly or not at all.

The relative locations of these three CBMs are quite far from each other, between 26 to 42 Å apart (Figure 4C). All three CBMs are in turn 30-36 Å away from the edge of the putative active site, and they fan out in different directions from it. The length of one arabinofuranose residue in an extended conformation is about 3.6 Å. This arrangement of the CBMs suggests that AftD may have evolved to bind long (>10 units) and possibly non-linear arabinan chains.

To test the above hypothesis, we probed the carbohydrate-binding ability of the protein *in vitro* by screening purified AftD against an array of mycobacterial glycan fragments (Zheng et al., 2017a). AftD appears to have specific binding to long and branched arabinofuranose chains (Figure 4D). Notably, the arabinofuranose glycans to which AftD binds have 10 or more arabinofuranose residues. Conversely, AftD appears not to bind to other mycobacterial glycans on the array: glycopeptidolipid, phosphatidyl*-myo-*inositol mannoside (PIM), phenolic glycolipids (PGL), trehalose mycolates and lipooligosaccharides (LOS), and capsular α glucans, with the exception of tetra-mannose which, surprisingly, appears to bind weakly (Figure 4D). These results corroborate both our structural hypothesis and the previously predicted function of AftD acting as a late arabinosyltransferase in the biosynthesis of LAM/AG (Alderwick et al., 2018; Škovierová et al., 2009).

### AftD is Complexed with Acyl Carrier Protein

At the C-terminus of the soluble domains of AftD, the protein chain re-enters into the membrane via TM 12, followed by an 18 residue loop that arches over TMs 5, 6, 7, 9 and 10 to reach TM 13 (Figures 2C and 5A). TMs 13-16 then form a four-helical bundle that interfaces with TM 8 and 9, which is unique to AftD in comparison to other Aft/Emb enzymes. This four-helical bundle shows a contiguous density adjoining the cytoplasmic side, consistent with the presence of an associated protein (Figure 5A). Using mass spectrometry, this additional component was identified as the acyl carrier protein (ACP) from *E. coli*, and the ACP crystal structure (PDB ID: 1T8K) was readily accommodated in the density (Figure 5A). The density map for ACP was of lower resolution (∼4-5 Å, Figure S3F) than for AftD, likely due to conformational flexibility. Indeed, focused classification of the ACP portion revealed multiple low-resolution conformational states. Notably, there were no classes without density for ACP, consistent with the tight association of the two proteins during expression and purification.

**Figure 5.**
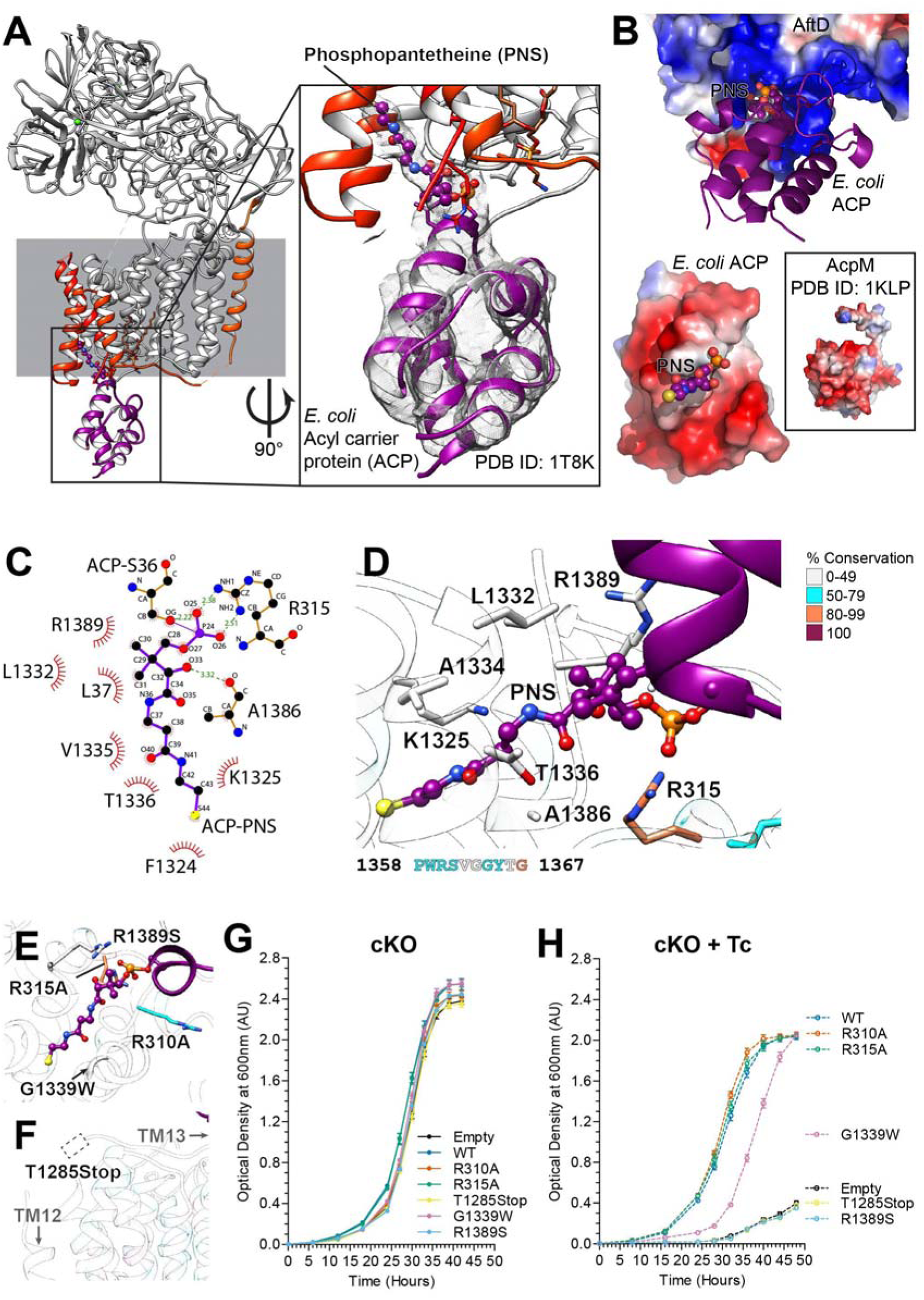
Acyl carrier protein (ACP) in complex with AftD. (A) TM 13, 14, 15 and 16 form a hydrophobic groove where the 4’-phosphopantetheine of ACP binds. AftD is rendered as cartoon with TMs 12-16 being colored in the same colors as in Figure 2. ACP is colored magenta. The insert shows a zoomed view of ACP and 4’-phosphopantetheine, with the corresponding density map shown as mesh. (B) Surface electrostatic potential maps of AftD (top), *E. coli* ACP (below) and *M. smegmatis* AcpM (below, insert), with red being negatively charge and blue positively charged. (C) LigPlot (Wallace et al., 1995) on 4’-phosphopantetheine showing its atomic interactions with AftD. (D) Zoom in on the binding interface between AftD and ACP, with side chains of relevant residues displayed as sticks. Residues have been colored by conservation using the same scheme as Figure 3 and 4. The primary sequence of the un-model flexible loop between TM15 and TM16 is shown below the insert, colored by conservation. (E) Residues that have been mutated in the *M. abscessus* AftD protein have side chains displayed as stick, and the model colored by conservation. If the residue is not resolved in the structure, its putative position is indicated by a dotted lined box (F).These mutations are also listed in a table (G). These mutant plasmids were then transformed into the cKO to assess their effect on the growth rate of *M. smegmatis* (H), in the absence or presence of tetracycline.

The negatively charged surface of *E. coli* ACP, which has an isoelectric point around 4, interacts with a positive patch of AftD comprising the arginine residues R310, R315 and R1389 (Figures 5B, 5C and 5D). ACP is found in its holo-form with 4’-phosphopantetheine covalently linked via a phosphodiester bond to S36 (Majerus et al., 1965). This prosthetic group can either be buried within the ACP hydrophobic tunnel-like cavity in the core of the four helices or extended towards its cognate protein (Cronan, 2014). The coulombic potential map shows that the 16 Å long 4’-phosphopantetheine fills the hydrophobic pocket of the four-helical bundle (TMs 13-16) of AftD (Figures 5C and 5D). Its C-terminal thiol group points towards the interspace between TM15 and TM16. Moreover, 4’-phosphopantetheine not only forms extensive hydrophobic interactions with AftD residues L1332, V1335 and F1324, but also establishes hydrogen bonds between its O33 and the backbone carbonyl of A1386, and its phosphate group with the side-chain of R315, a highly conserved residue among AftDs (Figure 5D).

To determine the physiological relevance of the ACP-AftD interaction, we purified *M. abscessus* AftD from a *M. smegmatis* mycobacterial expression system. Mass spectrometry analysis of the purified protein revealed the presence of *M. smegmatis* meromycolate extension acyl carrier protein AcpM (UniProtKB accession: ACPM_MYCS2, Table S2), a 99 residue protein that has 39% sequence identity with *E. coli* ACP. AcpM also has a negatively charged surface (PDB ID: 1KLP, Figure 5B insert) like the *E. coli* ACP (Wong et al., 2002). *M. smegmatis* has another ACP (UniProtKB accession: A0QUA2_MYCS2) which shares around 40% sequence identity with AcpM, and it was not detected via mass spectrometry. These results indicate that the presence of ACP in the AftD structure was not an artefact due to the overexpression system in *E. coli*.

Five mutants in the region of AftD that interacts with ACP were also generated with the aim of disrupting ACP binding. These were successfully purified in *E. coli* (Figure S6) and cloned into the mycobacterial expression vector. To investigate the functional relevance of the ACP-AftD interaction, we examined whether these mutant genes restored WT growth of the conditional *M. smegmatis aftD* mutant with tetracycline. The insertion of a stop codon at T1285 produced a C-terminal truncation that eliminated the four-helical ACP-binding bundle all together (Figure 5F). This truncated protein resulted in a growth defect akin to the empty plasmid, suggesting that either the four-helical bundle of the enzyme *per se*, or its interaction with ACP is important for AftD function (Figures 5G and 5H). Mutations were also made to three arginine residues (R310A, R315A, R1389S) to abolish (by mutations to alanine) or reduce (by mutations to serine) the charge interactions between the protein part of ACP and AftD (Figure 5E). R310A and R315A mutants fully restored growth of the conditional *M. smegmatis aftD* mutant in the presence of tetracycline, indicating these residues were not important for proper AftD function (Figures 5G and 5H). By contrast, R1389S showed a severe growth defect, consistent with a functional role of this residue (Figures 5G and 5H). Another mutation was introduced at G1339 to add a bulky tryptophan to prevent the binding of 4’-phosphopantetheine to AftD’s four-helical bundle (Figure 5E). This mutant only partially rescued growth of the *M. smegmatis aftD* cKO in the presence of tetracycline, suggesting that G1339 is also important for the function of AftD (Figures 5G and 5H).

To further characterize the two classes of AftD mutants that were made (targeting putative active site versus ACP binding), we performed microplate appearance (Figure 6A) and colony morphology assays (Figure 6B) on a representative mutant from each class: D474A (putative active site) and R1389S (ACP binding). Both mutants showed similar phenotypes to the cKO (Figures 1E and 1F) and to each other, suggesting that disruption of ACP binding may be linked to loss of enzymatic function.

**Figure 6.**
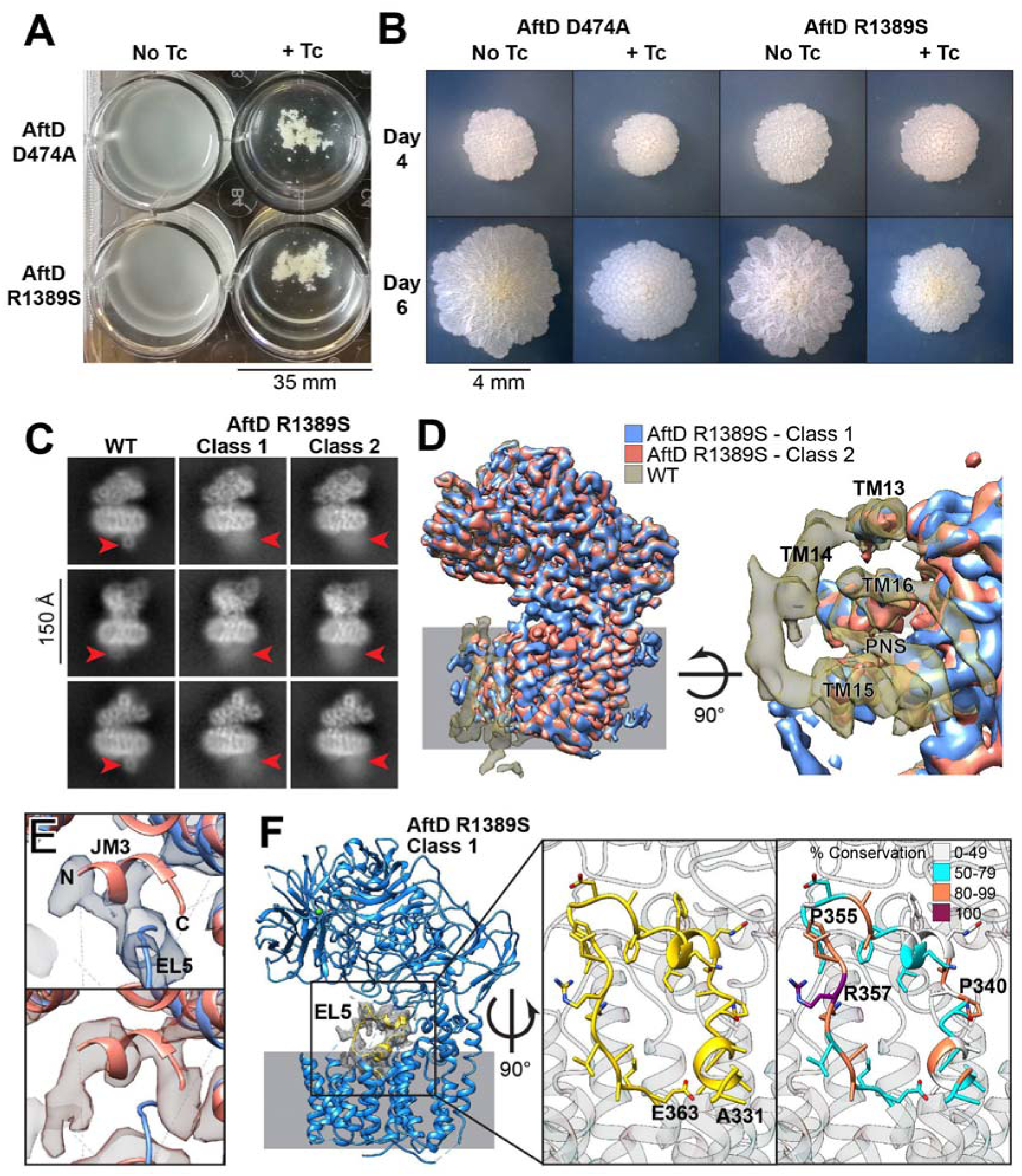
Phenotype and Structure of AftD-R1389S of *M. abscessus* AftD. (A) Microplate appearance assay and (B) colony forming assay across 4, 6 and 8 days of AftD mutants D474A and R1389S. (C) 2D class averages comparison between the AftD WT particles and mutant R1389S class 1 and 2 particles, with the red arrow pointing at the position of ACP. (D) Superposition of maps from class 1 and 2 from AftD mutant R1389S with the WT map. The maps have been low-pass filtered to 4 Å and displayed at similar thresholds for comparison. (E) Superposition of class 1 and 2 models from AftD-R1389S against class 1 map (on top) and class 2 map (on the bottom) at the JM3 region. Colored scheme follows (D). (F) The model of class 1 from AftD-R1389S with the ordered loop (EL5) colored in yellow. The rest of the structure is rendered as cartoon in blue. The inserts are zoomed around EL5 (residues 331 to 363), colored in yellow and by conservation.

### The Structure of AftD-R1389S Mutant Shows Ordering of EL5 Loop with Disordering of ACP

To investigate how ACP binding might be correlated to enzyme activity, we determined the structure of AftD-R1389S mutant using single-particle cryo-EM (Figures 6 and S7). The structure determination pipeline was similar to that described for wild-type AftD, with two notable exceptions: 1) TEV cleavage was not performed as the mutation resulted in TEV cleavage being very inefficient (Figures S7A and S7B); and 2) AftD-R1389S was vitrified on gold grids using the Leica GP, using conditions optimized earlier with Spotiton (Figure S7C). 4,886 micrographs were collected and 226,478 particles were initially processed following data collection on a Titan Krios equipped with a Gatan K2 Summit camera fitted with an energy filter (Table S1, Figures S7C and S7D). After data processing, the consensus reconstruction with an overall resolution of 3.4 Å was obtained from 150,978 final particles (Figure S7E). 2D class averages of the dataset revealed a fuzzy but identifiable density for the ACP – in comparison, the WT 2D class average showed a very distinct density with sharp features for the ACP (Figure 6C). This suggests that the mutation did not fully abrogate ACP binding, but caused it to be weakly associated with AftD-R1389S and hence highly flexible. This was corroborated by mass spectrometry analysis of the AftD bands from the SDS-PAGE gel (Figure S7B).

**Figure 7.**
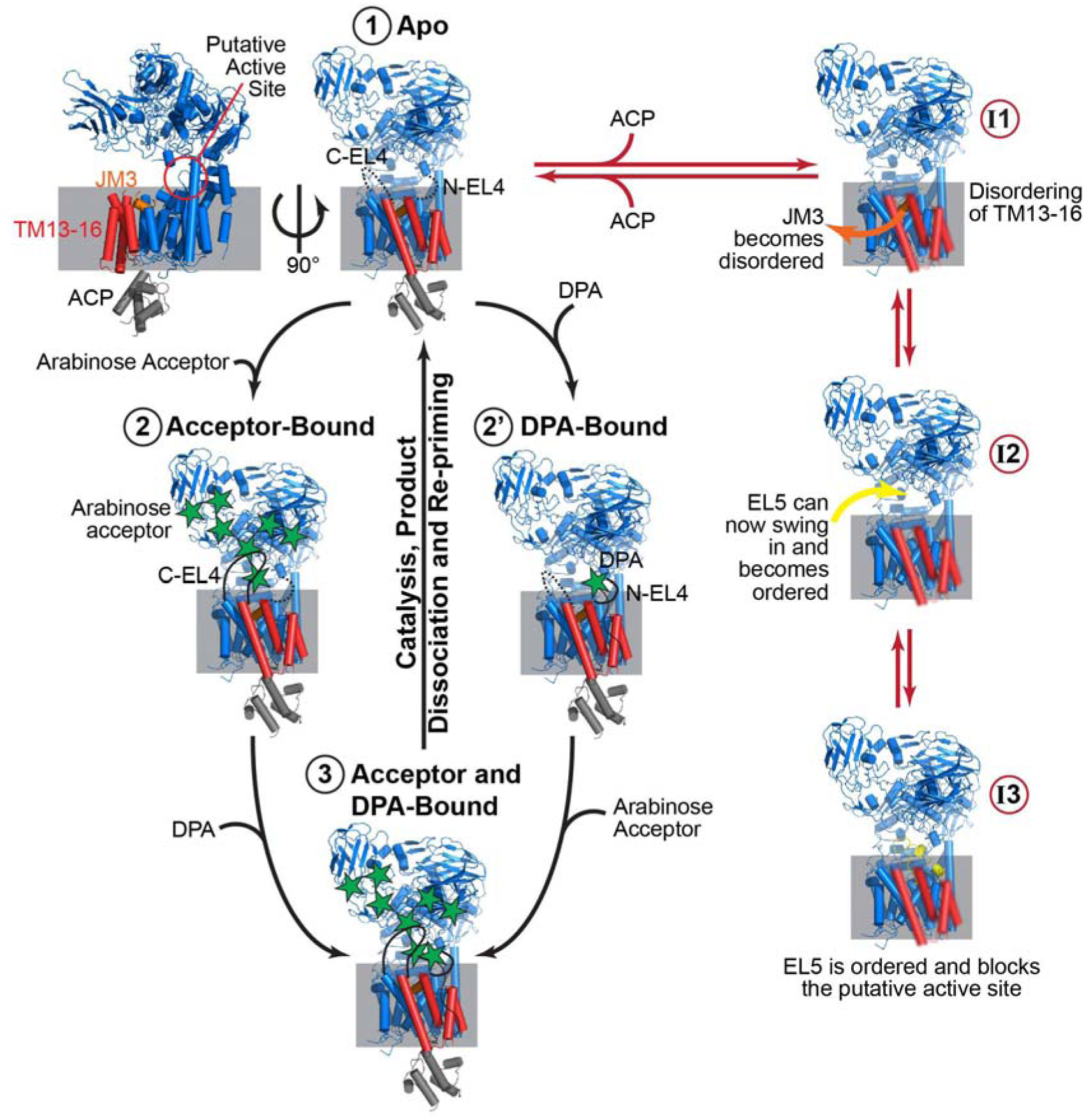
Postulated Mechanism of Action for AftD. AftD is rendered as cartoon in blue, orange (JM3), red (TM 13-16) and yellow (ordered EL5). ACP is rendered as cartoon in grey. The catalytic cycle for AftD is indicated by the black arrows from state 1 through 3. The inhibitory cycle is indicated by red arrows from state I1 to I3. The solved WT AftD structure corresponds to state 1, the AftD-R1389S class 2 structure to state I2 and the AftD-R1389S class 1 structure to state I3.

When inspecting the consensus reconstruction of AftD-R1389S, a distinct density near the active site that was not seen in the wild-type structure was observed (Figure S7E). Signal subtraction followed by focused classification was performed to resolve the heterogeneity, which produced two high resolution maps of AftD: class 1 at 3.5 Å and class 2 at 3.4 Å (Figures 6D, S7E, S7F and S7G). For both these classes, 2D class averages (Figure 6C) had fuzzy density for the ACP, and both of them also showed worse resolution for the last four TMs at ∼5 Å (Figure S7F). When the maps were low pass filtered to 4 Å and compared at similar display thresholds, TM14 was the most disordered helix and its density was hardly seen compared to the other helices (Figure 6D). The linker between TM14 and TM15 was also poorly ordered. No density was observed where the ACP bound to the WT structure, and no density for the 4’-phosphopantetheine was apparent in both AftD-R1389S classes (Figure 6D).

Between the two classes, the most notable difference was in the putative active site region (Figure 6F). Here, the class 2 structure highly resembles the WT structure. However, in class 1, there appears to be a significant conformational change. Firstly, the previously disordered EL5 loop between TM 9 and 10 is now resolved, and physically occludes the putative active site. Secondly, the previously disordered N- and C-EL4 loop between TM 7 and 8 still remain poorly ordered, but the JM3 now appears to be disordered with poor map density (Figure 6E). In comparison, JM3 is still present in class 2 (Figure 6E).

An atomic model was built into the resolved EL5 loop density, resulting in a U-shaped structure connecting TMs 9 and 10 (Figure 6F). Out of the residues present in the loop, R357 is absolutely conserved. Two prolines, P340 and P355, are responsible for the sharp turns in the loop structure, and both also have a high degree of conservation (85.1% and 93.1% respectively) (Figure 6F).

## DISCUSSION

### The Conditional *aftD* Mutant in *M. smegmatis* Provides a System to Examine Structure-Function Relationships of AftD

In this study, we constructed a tetracycline-inducible conditional *aftD* mutant in *M. smegmatis* and showed that AftD is essential in *M. smegmatis* as reported previously (Škovierová et al., 2009) in contrast to closely related corynebacteria which produce a much shorter AftD protein (Figure S2A) (Alderwick et al., 2018). We observed that AftD depletion increased cell clumping and altered the colony morphology of *M. smegmatis* (Figure 1). Both phenotypes are consistent with a change of the cell surface properties and the function of AftD in adding to long, branched arabinan as shown in this study (Figure 4D). The branched arabinan chain forms an integral part of the arabinogalactan polymer and of the attached outer membrane (Hoffmann et al., 2008), which is essential in mycobacteria, but not in corynebacteria (Portevin et al., 2004). Presumably, interfering with the arabinogalactan synthesis disturbs the proper assembly of the outer membrane and its properties, thus explaining the observed phenotypes. AftD depletion resulted in shorter *M. smegmatis* cells in contrast to the phenotype of a previous report which utilized a temperature-sensitive rescue plasmid to deplete AftD from *M. smegmatis* at 42 °C (Škovierová et al., 2009). This discrepancy might result from the non-physiological temperature in the latter experiment which is known to trigger a heat shock response in mycobacteria altering expression of many genes (Stewart et al., 2002) and protein properties. By contrast, our conditional mutant enables for the first time to examine the function of AftD in mycobacteria under physiological conditions.

### The Full-Length Structure of a Mycobacterial Membrane Glycosyltransferase

The GT-C membrane glycosyltransferase family (Liu and Mushegian, 2003) is underrepresented in the protein databank (PDB), with currently 13 structures of just five different GT-C members (Bai et al., 2019; Bai et al., 2018; Matsumoto et al., 2013; Matsumoto et al., 2017; Napiórkowska et al., 2018; Napiórkowska et al., 2017; Petrou et al., 2016; Wild et al., 2018). None of these previous structures are of mycobacterial GT-C glycosyltransferases. The AftD structure presented herein is the first full-length structure of this family, and the first determined by cryo-EM (Figure 2). The high degree of structural conservation of AftD with other GT-C structures across all three domains of life indicates a remarkable retention of the fold across evolutionary time and also points to the likely existence of a common ancestral GT-C fold (Figure 4C). However, the structural conservation is not uniform – the TM regions are marginally better conserved than the soluble GT-C domain, an observation also predicted previously (Lairson et al., 2008). The variability of the soluble domain is likely due to variations in substrates of each GT-C: for the four GT-C members where atomic structures are available, their substrates are either lipid-linked glycans and peptides (STT3, PglB, AglB and Pmt1-Pmt2), or lipid-linked glycans and lipid A (ArnT). AftD differs from these as its acceptor is thought to be a complex glycan (Figure 4D), providing an explanation for the fact that it has three additional CBM domains attached to its soluble GT-C domain.

### Arabinosyltransferase Function of AftD

Structure-based alignment with the other GT-C glycosyltransferases of known structure, allowed us to locate the putative active site of AftD in a cavity between the membrane-bound TM and soluble GT-C domain (Figure 2). We confirmed, by mutagenesis studies, that five of the conserved residues (K39, D41, E251, K361, D474) around the putative active site are essential for function and we thus postulate that these are involved in either catalysis or substrate binding. For substrate binding, these residues need to interact either with the phospho-arabinofuranose moiety of DPA, terminal arabinofuranose(s) of arabinogalactan/lipoarabinomannan, or the divalent cation likely required for catalysis (Sharma et al., 1981). The map shows poorly resolved densities for the flexible EL4 loop around this site for apo-AftD, and we expect this region to become ordered to assist in substrate binding, as shown to occur in PglB and ArnT (Napiórkowska et al., 2017; Petrou et al., 2016). N-EL4 should become ordered to trap DPA (State 2’ in Figure 7), being closest to the putative DPA binding site (Figures S5B, S5D, S5E and S5F). On the other hand, C-EL4 is well-positioned to become ordered when the growing end of the arabinan acceptor enters the active site (State 2 in Figure 7). In our structure, no divalent cation density around the putative active site was observed. In analogy with PglB, we predict that substrate binding and ordering of the EL4 loop might be required to trap a catalytic cation (Napiórkowska et al., 2017).

We identified three putative CBMs in the periplasmic domain, and showed the possible glycan binding interfaces and residues by comparison to other structurally homologous CBMs for which ligand-bound structures are available (Figure 4). The sizeable distances between the AftD CBMs (26 to 42 Å, Figure 4C) suggests the binding of a large, complex glycan substrate, which corroborates previous work placing AftD as a late stage glycosyltransferase that acts near the end of the synthesis pathway for the arabinans in both LAM and AG, (Alderwick et al., 2018; Škovierová et al., 2009). As observed previously in nature, an example being cellulase Cel5A (Boraston et al., 2003), which has two CBMs, CBM multivalency carries two advantages: 1) modules work together cooperatively to result in stronger binding than individual CBMs, and 2) each module recognizes different regions of the substrate, thus improving binding specificity. In AftD, these domains, which rise 70 Å up from the membrane (Figure 2A), are likely to bind specifically to different portions of AG/LAM. The glycan array result supports this hypothesis: branched arabinofuranose glycans >10 residues in length, which are too large to be bound by a single CBM, appear to show the most convincing binding profiles (Figure 4D). The fact that both elongating AG and LAM are restrained in one dimension, being anchored via a polyprenol-pyrophosphate in the periplasmic membrane (Alderwick et al., 2015), and to either the periplasmic membrane through phosphoinositol mannoside (Kordulakova et al., 2002; Lea-Smith et al., 2008; Morita et al., 2006) or possibly also to the outer mycolate membrane (Alsteens et al., 2008), respectively, will likely significantly impact binding affinities.

Combining our results with the knowledge derived from other GT-C membrane glycosyltransferases (Napiórkowska et al., 2017; Petrou et al., 2016) studied both structurally and functionally, we propose the following putative enzymatic cycle (Figure 7). The solved AftD structure represents the apo-state, bearing disordered N-EL4 and C-EL4 regions (State 1 in Figure 7). Binding of the substrates will result in ordering of N-EL4 and C-EL4, respectively, stabilization of the substrate-enzyme complex (State 2 or 2’ in Figure 7), and priming of the system for catalysis to occur (State 3 in Figure 7). Thereafter the product is released and the active site is re-primed. This would reset AftD back to the apo-state, readying it for another catalytic cycle.

Further understanding of the order of substrate binding and of the precise structural rearrangements that occur in the active site will require structures of substrate bound of AftD. Attempts to obtain such structures have been made – we have tried adding various glycan moieties as indicated from our glycan array results, decaprenyl-phosphate and an analog of DPA (Joe and Lowary, 2006) to obtain a substrate bound conformation of AftD. No ligand density was observed in these preliminary structures, possibly due to the transient nature of binding of such natural ligands, technical difficulties in substrate incorporation, or presence of the *E. coli* (non-native) ACP that might be inhibitory. An alternative strategy might involve the chemical synthesis of substrate analogues or inhibitors of AftD that have higher binding affinity than the endogenous counterparts.

Such an approach has previously yielded positive results for PglB (Napiórkowska et al., 2017) and STT3 (Bai et al., 2018; Wild et al., 2018).

### Role of the Acyl Carrier Protein in AftD

In the structure of AftD, an *E. coli* ACP was so tightly bound to the cytoplasmic face of the enzyme at a C-terminal four-helical bundle, that it remained associated during purification (Figure 5A). The tight binding of the ACP is quite remarkable because complexes of ACP with partner proteins have typically been difficult to observe due to the transient and weak nature of their interactions (Cronan, 2014). Indeed, some of these complexes could only be obtained via crosslinking experiments (Tallorin et al., 2016). Neither the ACP nor the four-helical bundle it associates with is present in other GT-C glycosyltransferases, for which structural information is available. This interaction was found to be: 1) physiological, as AftD purified from *M. smegmatis* also remained associated with the *M. smegmatis* ACP homolog AcpM (Table S2), and 2) likely essential for function, as AftD variants genetically engineered to disrupt AcpM binding did not rescue the growth defect of AftD-depleted *M. smegmatis* (Figure 5H).

Structurally, mycobacterial ACP is very similar to its *E. coli* counterpart, with helix II (the recognition helix) being the most conserved (Wong et al., 2002). The only notable difference is the presence of a ∼35 residue extension at the C-terminus, which is highly flexible and is thought to interact with long chain intermediates of mycolic biosynthesis or mediate specific protein-protein interactions (Figure 5B) (Schaeffer et al., 2001). The highly conserved helix II contains the serine 36 residue, which is located within a signature motif of D<u>S</u>L/X-hydrophobic residue and is modified by a 4’-phosphopantetheine that interacts with AftD in our structure. The high degree of conservation of ACP across species is the likely reason why we were able to isolate an ACP-AftD complex from a heterologous expression system. In contrast to *E. coli*, the genome of *M. smegmatis* encodes two different ACPs. The AcpM variant is involved in both fatty acid (Kremer et al., 2001) and mycolic acid synthesis (Zimhony et al., 2015).

As no further electron density was observed beyond the terminal thiol group of the 4’-phosphopantetheine, it is not clear whether the ACP has no cargo bound, a disordered cargo, or a variety of different cargos in different conformations that average out during cryo-EM analysis. In AftD, the ACP could be serving as a shuttle for starting materials and intermediates throughout the fatty acid biosynthetic pathway. In *E. coli*, ACP interacts with at least twelve enzymes involved in fatty acid biosynthesis, as well as with seven other enzymes from disparate biosynthetic pathways (Nguyen et al., 2014). In mycobacteria, the ACP also functions as a shuttle for mycolic acid biosynthesis (Takayama et al., 2005). Intriguingly, ACP has been reported to be essential for an *E. coli* enzyme catalyzing the synthesis of β-(1→2) -linked glucan in membrane-derived oligosaccharides from UDP-glucose, although the 4’-phosphopantetheine moiety was not needed (Therisod and Kennedy, 1987).

The elucidation of the structures of mutant AftD-R1389S (Figures 6 and S7) pointed to another hypothesis for the presence of the ACP – as a possible regulator of AftD enzyme activity (Figure 7). ACP regulation of catalytic activity has indeed been observed in other proteins like WaaP (Kreamer et al., 2018) and possibly SpoT-like proteins (Battesti and Bouveret, 2009). When the 4’-phosphopantetheine is removed from the four-helical bundle, the helices become more disordered, as evidenced by the lower resolution, especially for TM14 (Figure 6D and State I1 in Figure 7). This, in turn, results in JM3 moving away and the EL5 loop becoming ordered (Figure 6E and State I2 in Figure 7). The ordered EL5 now appears to block the putative active site, possibly preventing substrates from entering, and thus effectively halting enzymatic function (Figure 6F and State I3 in Figure 7). Out of the two solved dominant structures of AftD-R1389S, class 2 may represent an intermediate state in this conformational change as the four-helical bundle is disordered but JM3 is still present. Class 1, on the other hand, likely represents the final state of this postulated conformational change, with EL5 ordered and JM3 displaced. The fact that a mutation of the enzyme active site (AftD-D474A) results in *M. smegmatis* that looks phenotypically similar to that of AftD-R1389S is consistent with the observation that both these mutations could abrogate the same function of AftD – presumably its glycosyltransferase activity (Figures 6A and 6B). As the ACP binds the cytoplasmic face of AftD, this could enable the transmission of intracellular signals across the membrane. Given that AftD is the crucial arabinofuranosyltransferase that adds the final α-linked arabinofuranose sugars on AG (Alderwick et al., 2018), inactivating this enzyme could effectively prevent the attachment of mycolic acids and block the proper assembly of the entire outer membrane.

### AftD as a Potential Drug Target

AftD is an essential gene in mycobacteria and is found on the inner membrane (Figure 1A), making it an attractive target for the development of novel anti-tubercular antibiotics (Favrot and Ronning, 2012). There are no current drugs in the market that are known to target AftD, although there have been reports of mutations on the *aftD* gene that are associated with drug resistance and susceptibility. The T797A mutation in *M. tuberculosis* AftD results in resistance to isoniazid (Jena et al., 2016; Shekar et al., 2014), whereas S1080G in *M. tuberculosis* AftD results in multiple drug resistant forms of the pathogen (MDR and XDR) to become susceptible to amoxicillin and clavulanic acid (Cohen et al., 2016). Both T797 and S1080 are on poorly conserved regions on the surface of the soluble region of AftD, making their functional or structural relevance difficult to interpret.

Two additional factors make AftD an attractive potential drug target. Firstly, AftD is well conserved across many other bacteria of the *Actinobacteria* class, which includes other pathogens that cause diseases like diphtheria (*Corynebacterium diphtheria*), leprosy (*M. leprae* or *M. lepromatosis*) and nocardiosis (various *Nocardia* species). Secondly, the fact that AftD has two potential sites at which to target drugs (the active site and the ACP binding site) that both appear to be essential for function, further contributes to the potential of this protein for structure-based drug design efforts.

## Conclusions

Here, we present the first full-length mycobacterial GT-C structure, determined by single-particle cryo-EM to 2.9 Å resolution. The structure of wild-type AftD reveals a conserved GT-C fold and also pinpoints the location of the putative active site. The residues that are likely involved in substrate binding and catalysis were probed through mutagenesis studies using an engineered AftD cKO strain in *M. smegmatis*. Furthermore, the structure and glycan array analysis show that AftD binds specifically to large, highly branched arabinofuranose residues that are found in AG/LAM. Finally, to the best of our knowledge this is the first report of a glycosyltransferase structure with a tightly associated ACP. This interaction, when disrupted using a specific single mutant (AftD-R1389S), resulted in a series of conformational changes in AftD, as observed in the structures of this mutant. These changes lead to ordering of a loop that blocks the putative active site and could be a mechanism for transmitting the signal from the cytoplasmic face of AftD to the periplasmic active site.

## AUTHOR CONTRIBUTIONS

F.M. & O.B.C. conceived the study. A.L.R., B.K., and J.R. performed the genomics expansion and small scale screening. J.R. did medium scale expression and purification. Y.Z.T. did the large scale expression, purification and negative stain EM. V.P.D., H.W. and Y.Z.T. vitrified the grids. Y.Z.T. collected, processed and analyzed the cryo-EM data. Y.Z.T. built the model with O.B.C’s help. Y.Z.T. did the phylogenetic analysis. L.Z. constructed and characterized the conditional *aftD* deletion mutant in *M. smegmatis* and analyzed the function of AftD mutants in *M. smegmatis*, with M.N.’s supervision. A.M.R. and Y.Z.T. collected the scanning electron micrographs. L.Z., Y.Z.T. and S.L.G. constructed the AftD mutants. R.B.Z. did the glycan array experiments with T.L.L.’s supervision. J.R. and D.A. pulled down AftD from *M. smegmatis*, with advice from M.J.C. and M.P., under M.A.’s supervision. B.C. & C.P. supervised EM analysis. F.M. supervised the entire project. Y.Z.T. and F.M. wrote the manuscript, with input from all authors.

## ACKNOWLEDGEMENTS

We thank Joel Nott (Iowa State University) and Ricardo Gomes (UniMS, ITQB/IBET, Oeiras) for protein mass spectrometry help. We thank Ed Eng, Bill Rice, Laura Kim, Mikhail Kopylov and Kelsey Jordan (New York Structural Biology Center, Simons Electron Microscopy Center) for help with microscope setup. We thank Sargis Dallakyan, Carl Negro, Shaker Krit and Swapnil Bhatkar (New York Structural Biology Center, Simons Electron Microscopy Center) for computation support. We thank Dirk Schnappinger (Weill Cornell Medicine) for providing the components of the reverse TetR system. We thank Jeremie Vendome (Schrödinger) and Dmitry Lyumkis (Salk Institute) for critical reading of the manuscript. We thank Christina Chen for help with nanodisc incorporation, Khuram Ashraf for help with expression optimization, Vasileios Petrou for helpful discussions and Leora Hamberger for her assistance managing the Mancia laboratory (Columbia University). This work was supported by grants from the NIH (P41 GM103310 to C.S.P., B.C..; R01 GM111980, R35 GM132120 and R21 AI119672 to F.M.), the Agency for Science, Technology and Research Singapore (to Y.Z.T.), the University of Alabama at Birmingham (to M.N.), the Fundação para a Ciência e Tecnologia, Portugal (PD/BD/128261/2016 to J.R.; PTDC/BIA-BQM/30421/2017 and IF/00656/2014 to M.A.; PTDC/BIA-MIC/31233/2017 to M.J.C.), the Simons Foundation (SF349247 to C.S.P., B.C.), NYSTAR (to C.S.P., B.C.) and the Alberta Glycomics Centre (to T.L.L.). Some of the work was performed at the Center for Membrane Protein Production and Analysis (COMPPÅ; P41 GM116799 to Wayne Hendrickson) and at the National Resource for Automated Molecular Microscopy at the Simons Electron Microscopy Center (P41 GM103310), both located at the New York Structural Biology Center. M.A. acknowledges MostMicro Research Unit (financially supported by LISBOA-01-0145-FEDER-007660 funded by FEDER funds through COMPETE2020 and by national funds through FCT), and iNOVA4Health (LISBOA-01-0145-FEDER-007344, co-funded by FEDER under PT2020).

---

**Figure S1.**
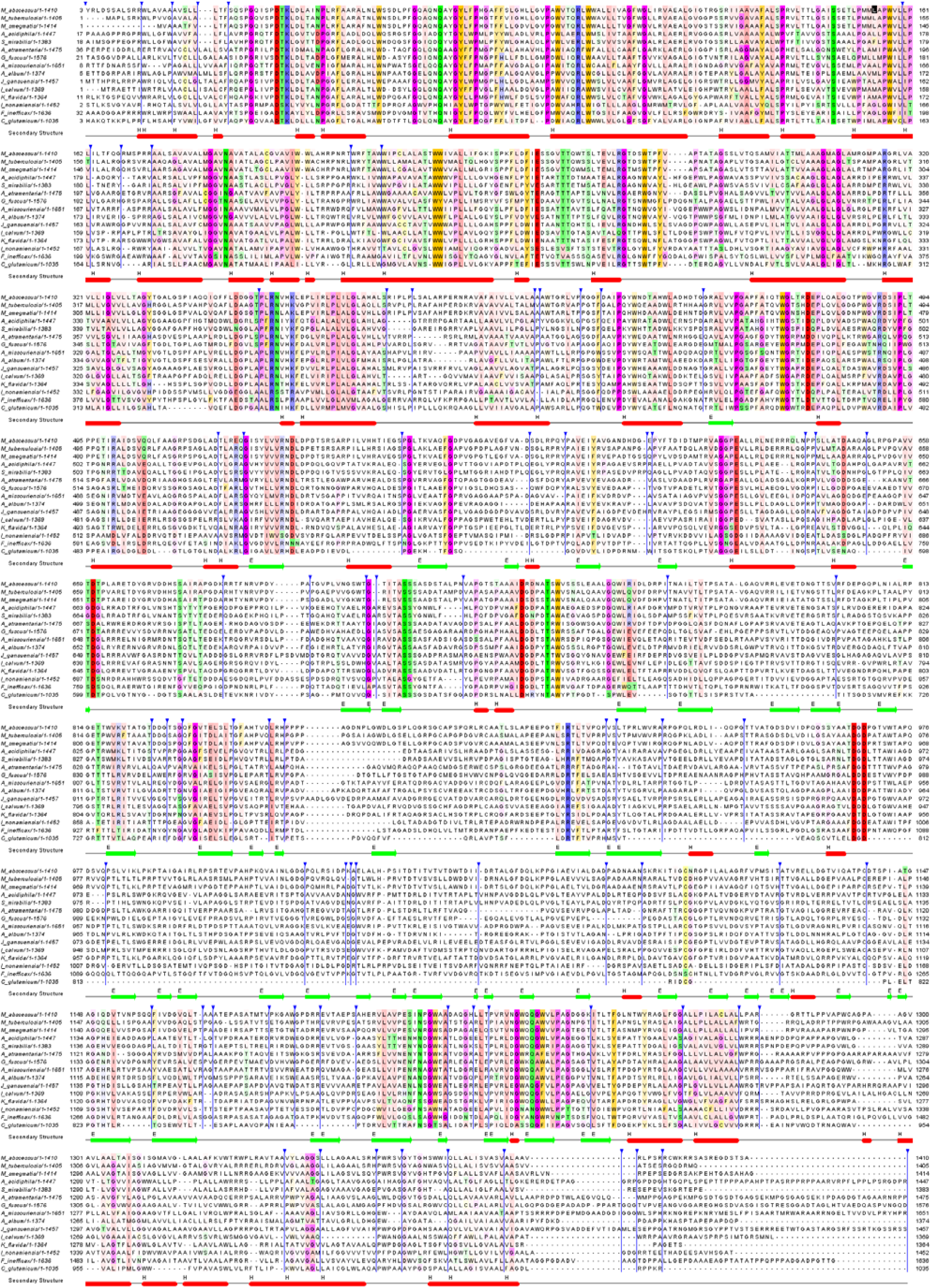
Multiple sequence alignment of AftD across the *Actinobacteria* class, Related to Figure 1. The sequence alignment was generated using MUSCLE and comprises of the following species (order in brackets): *Mycobacterium abscessus (Corynebacteriales), Mycobacterium tuberculosis (Corynebacteriales), Mycobacterium smegmatis (Corynebacteriales), Actinospica acidiphila (Catenulisporales), Streptomyces mirabilis (Streptomycetales), Actinomadura atramentaria (Streptosporangiales), Glycomyces fuscus (Glycomycetales), Actinoplanes missouriensis (Micromonosporales), Alloactinosynnema album (Pseudonocardiales), Jiangella gansuensis (Jiangellales), Intrasporangium calvum (Micrococcales), Kribbella flavida (Propionibacteriales), Ilumatobacter nonamiensis (Acidimicrobiales), Frankia inefficax (Frankiales)* and *Corynebacterium glutamicum (Corynebacteriales).* The residues are colored by their physiochemical properties using the Zappo scheme: Aliphatic/hydrophobic residues are in salmon (ILVAM), aromatic residues in orange (FWY), positively charged residues in blue (KRH), negatively charged residues in red (DE), hydrophilic residues in green (STNQ), conformationally special residues in pink (PG) and cysteines in yellow (C). The intensity of the coloring is proportional to the degree of conservation. Secondary structure elements derived from the *M. abscessus* structure are labelled and aligned below. Gaps and unaligned parts of the sequence have been shrunk and indicated by the blue arrow.

**Figure S2.**
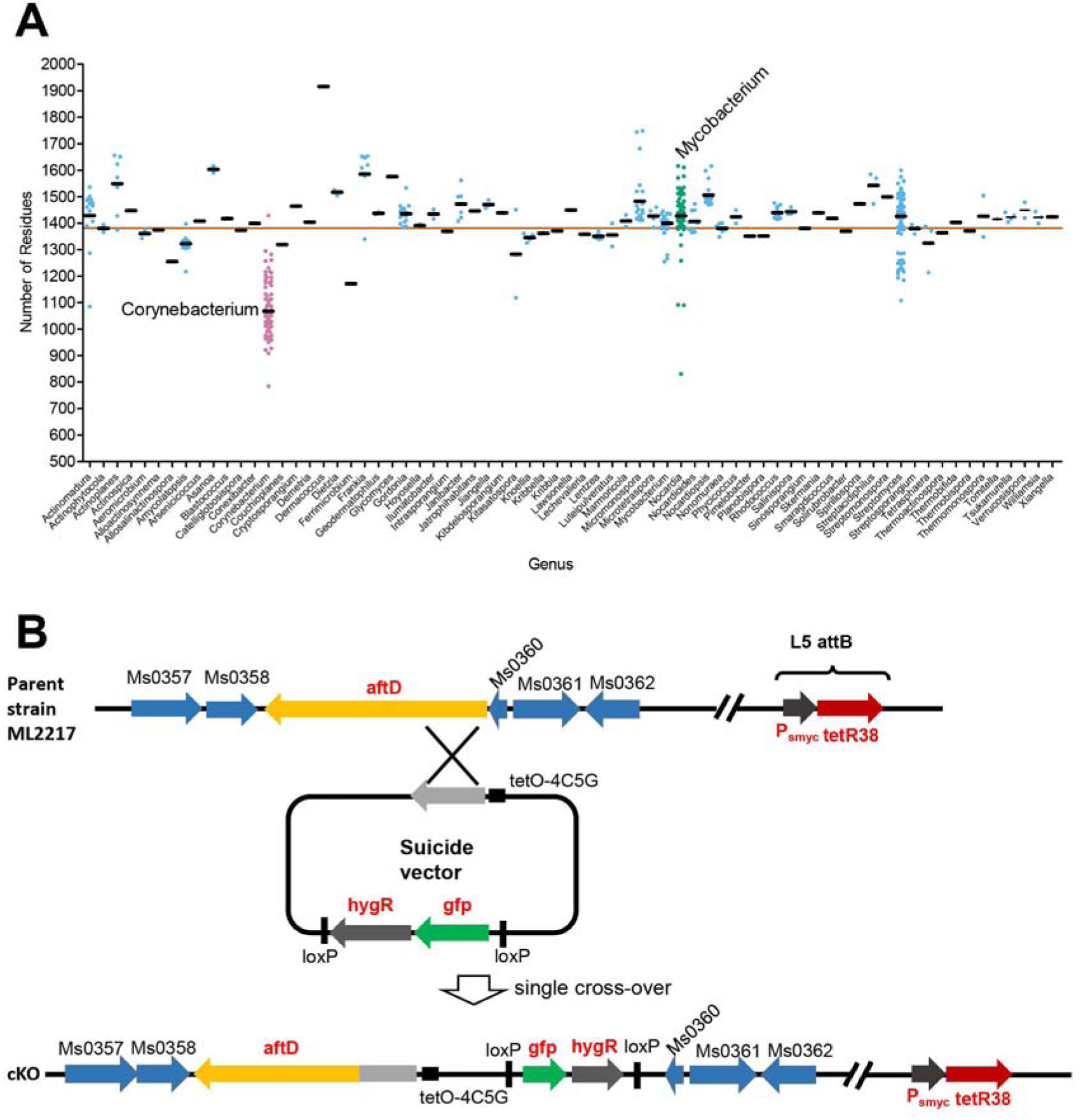
Lengths of AftD Homologs and Conditional Knock-out Scheme, Related to Figure 1. Total number of residues of each AftD homolog against the genus. Corynebacterium genus (pink dots) has unusually short AftD sequence compared to the rest of the genus in the *Actinobacteria* class (blue dots), including the *Mycobacterium* genus (green dots). (B) Conditional knock-out scheme for *aftD* gene in *M. smegmatis*. The gene locus for WT *M. smegmatis* around the *aftD* gene is shown.

**Figure S3.**
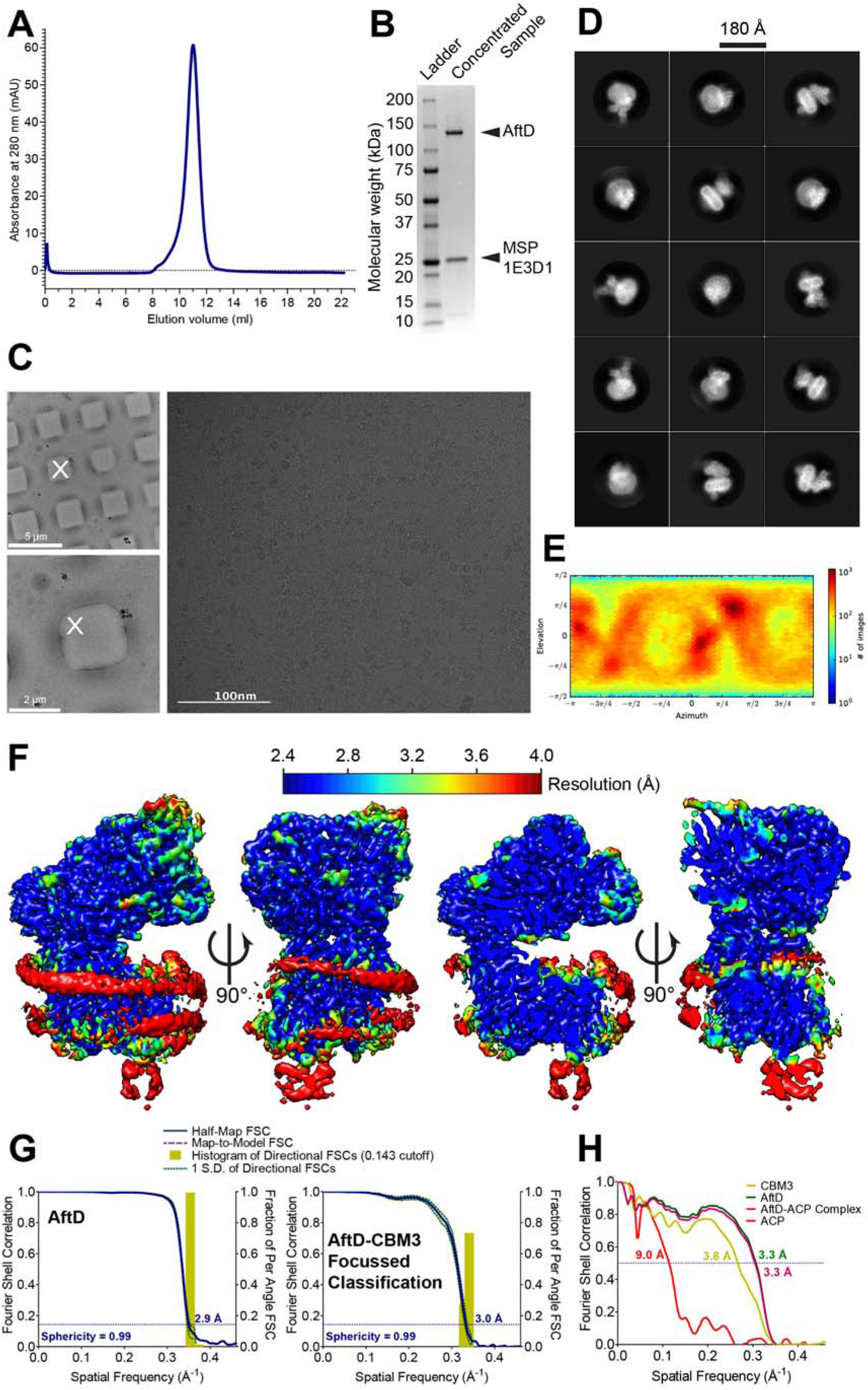
Purification and single-particle cryo-EM structural determination of wild-type M. abscessus AftD, Related to Figure 2. (A) Representative size-exclusion chromatography (SEC) trace of AftD incorporated into MSP-1E3D1 nanodiscs. Fractions corresponding to the peak were used for cryo-EM analysis. (B) SDS-PAGE gel of the concentrated sample from the peak fractions stained with Coomassie blue. The predicted molecular weight of AftD is 150 kDa, while that for MSP-1E3D1 is 32 kDa. (C) Representative square, hole and exposure magnification views of the Spotiton nanowire carbon patterned grids that were imaged. The exposure image had a defocus of -1.7 μ (estimated by CTFFind4 and pixel size of 1.0605 Å/pixel. (D) Representative 2D class averages. (E) Euler angle distribution plot of the final 3D reconstruction from CryoSPARC 2. (F) ResMap display of unsharpen AftD reconstructions in orthogonal views, showing the nanodisc belt as well as variation in local resolution. (G) Fourier shell correlation (FSC) curves for the full AftD map and AftD-CBM3 focused classified map describing the half-map (blue) resolutions (at 0.143 cut-offs), as well as histogram of directional resolutions sampled evenly over the 3DFSC (yellow). The corresponding sphericity value is also indicated. (H) Fourier shell correlation (FSC) curves for the atomic models of AftD, ACP, CBM3 of AftD and AftD with ACP against the whole composite map generated by combining the full AftD map with the AftD-CBM3 focused classified map using phenix.combine_focused_maps at 0.5 cut-off.

**Figure S4.**
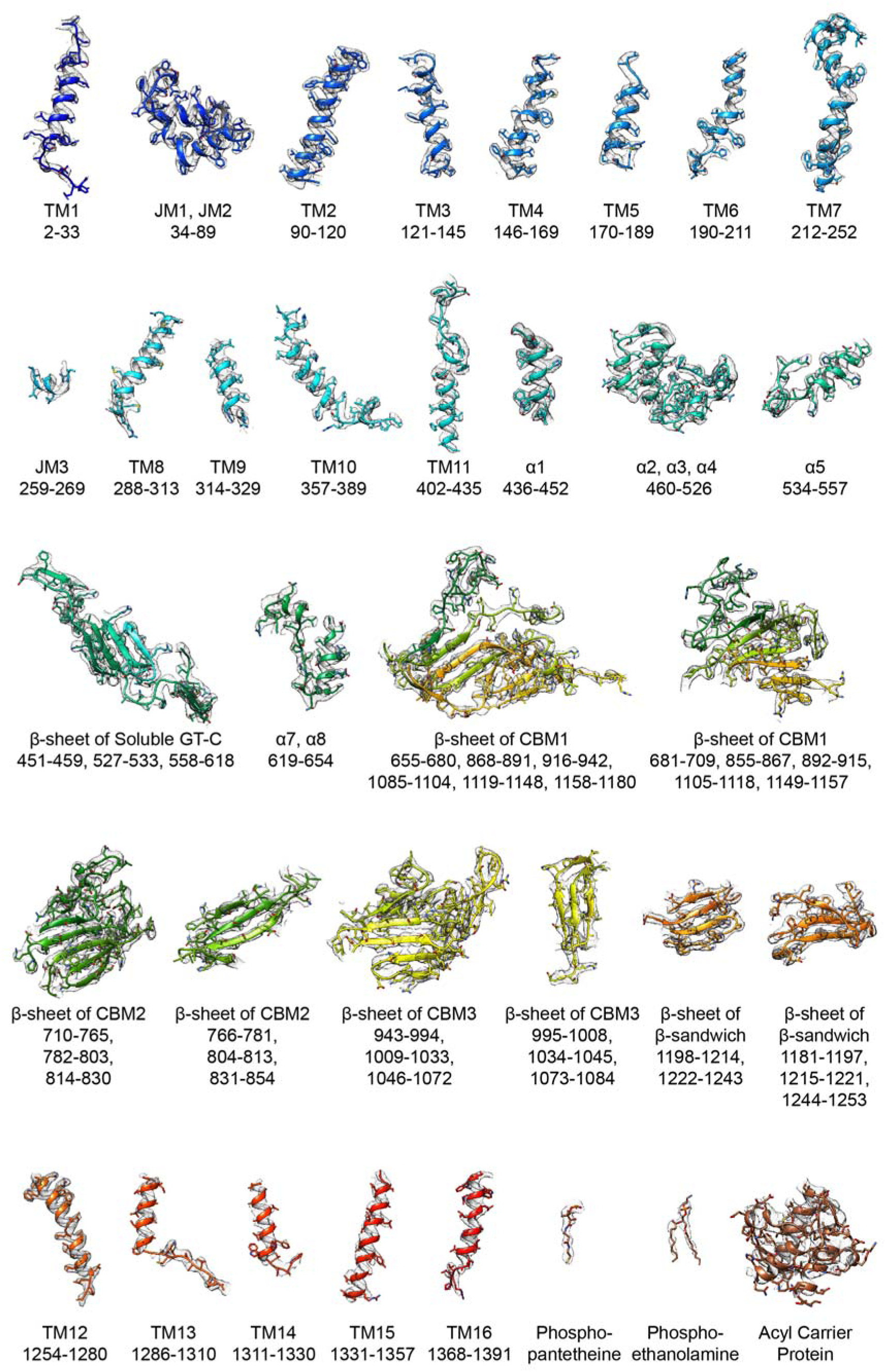
EM Density of AftD, Related to Figure 2. The atomic model for the structure of wild-type *M. abscessus* AftD is colored in rainbow and rendered as cartoon, with the side chains rendered as sticks. The map density is displayed as a mesh. The residues for each segment of the atomic model are indicated below the name of the segment. For all the segments except the acyl carrier protein, the sigma value for display was 8.33, while for the acyl carrier protein, it was 2.44.

**Figure S5.**
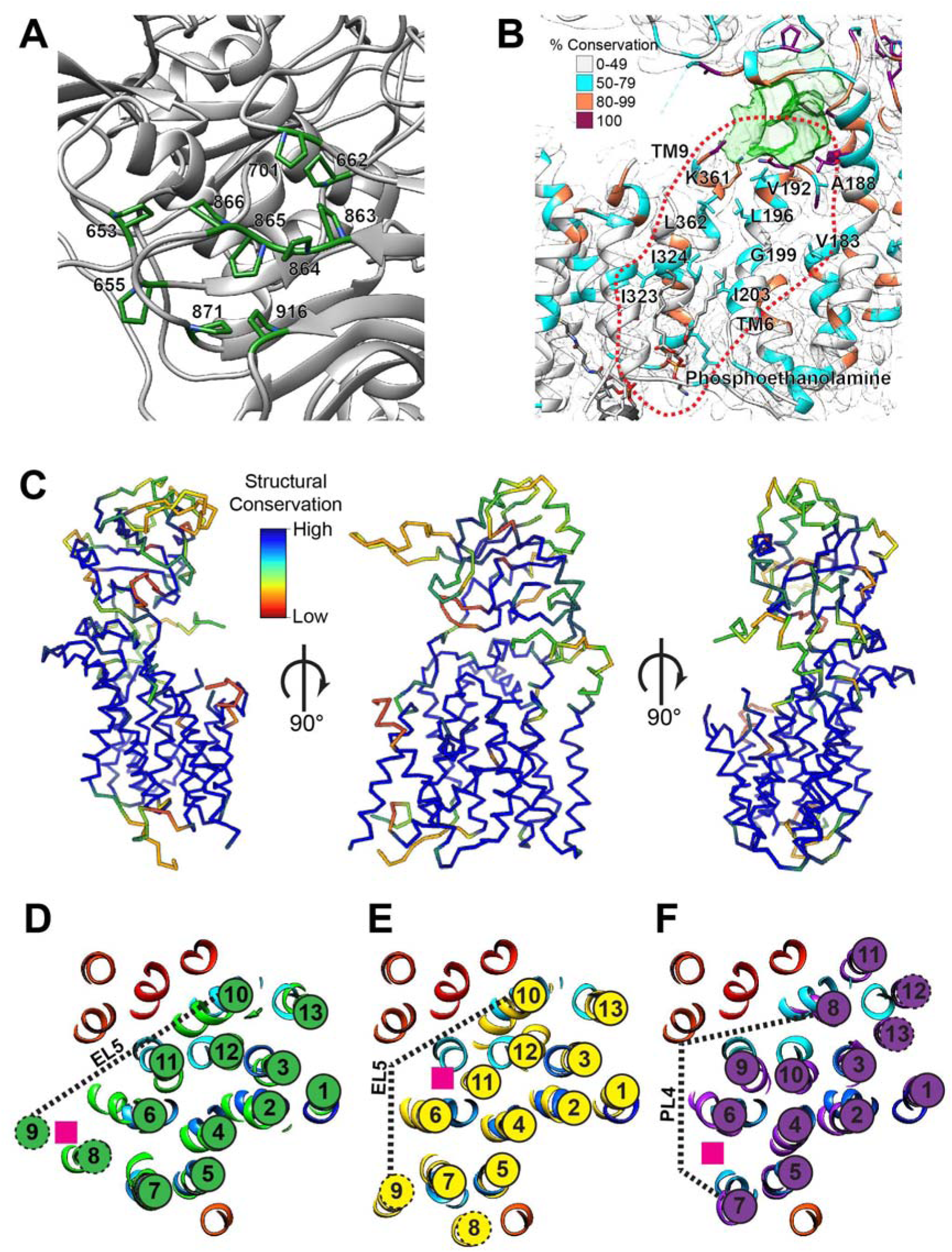
Structural Features of AftD, Related to Figures 2,3,4,5. (A) Stretch of consecutive prolines in CBM2 and neighboring prolines shown as green sticks. The rest of the structure is rendered as cartoon. (B) Location of the potential binding site (dotted red line) for the lipidic tail of the donor DPA. AftD is rendered as cartoon, with residues colored by conservation. The density map is shown as transparent grey. (C) Structural conservation of the GT-C fold of AftD (residues 1 to 600) against other GT-C glycosyltransferases obtained from a Dali search. Thereafter, (D) eukaryotic STT3 (PDB ID: 6EZN), (E) bacterial PglB (PDB ID: 5OGL) and (F) ArnT (PDB ID: 5EZM) were then superimposed onto full-length AftD and a slice through the transmembrane helices are shown as cartoon. The other glycosyltransferase structures were colored in green, yellow and purple respectively, while full-length AftD was colored in rainbow. Helices of the other glycosyltransferases were labelled, and those that did not have a corresponding homologous helix in AftD were indicated with a dotted circle. Locations of the flexible loop that becomes ordered upon substrate binding (EL5, PL4) are indicated by the dotted lines. The locations of the lipidic sugar donor substrate are indicated by pink boxes.

**Figure S6.**
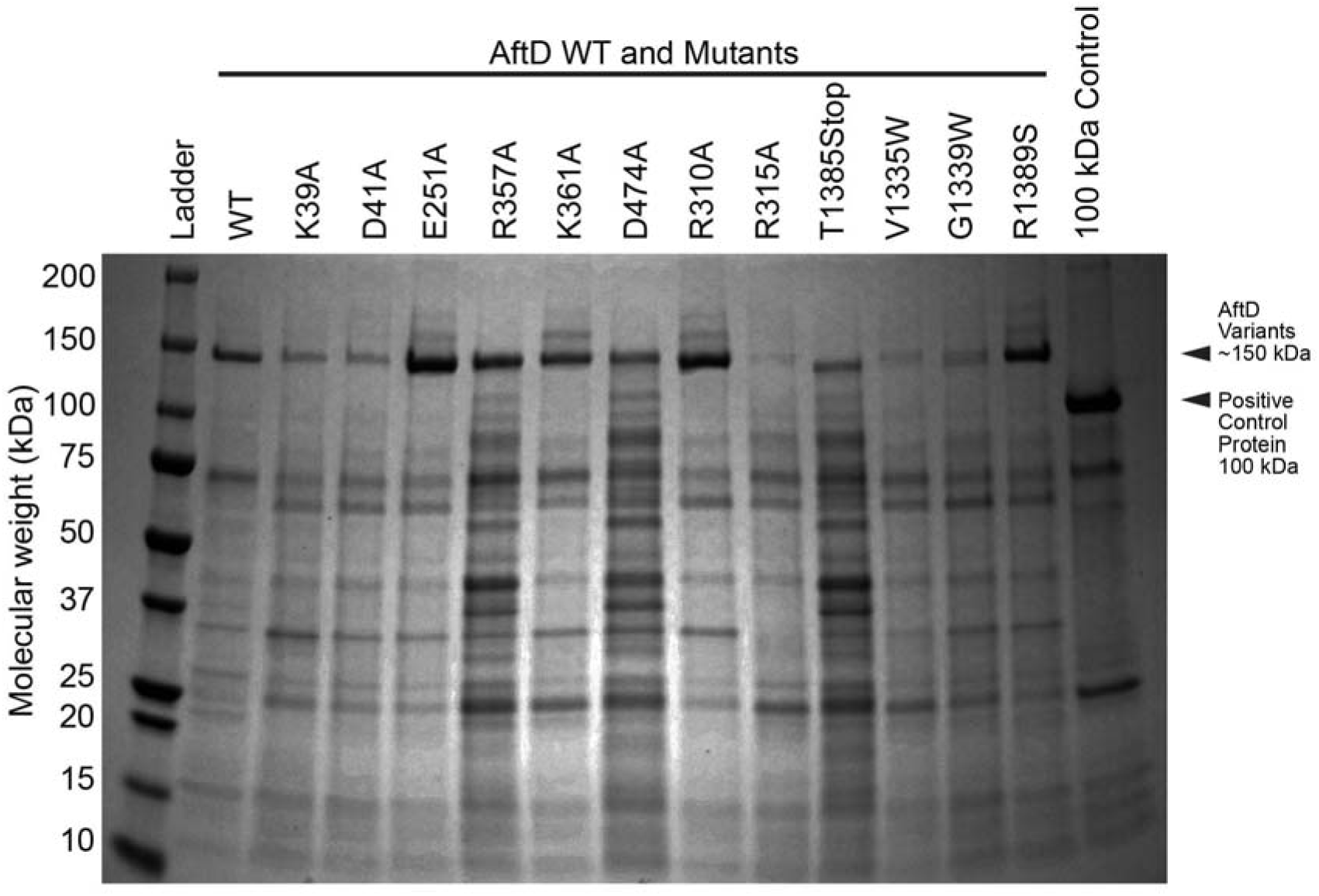
Expression of AftD mutants in *E. coli*, Related to Figures 3,5. SDS-PAGE gel of samples from small scale purifications of WT AftD and AftD mutants, stained with Coomassie blue. The AftD WT and mutants were heterologously expressed in *E. coli* and purified for this experiment, without TEV cleavage.

**Figure S7.**
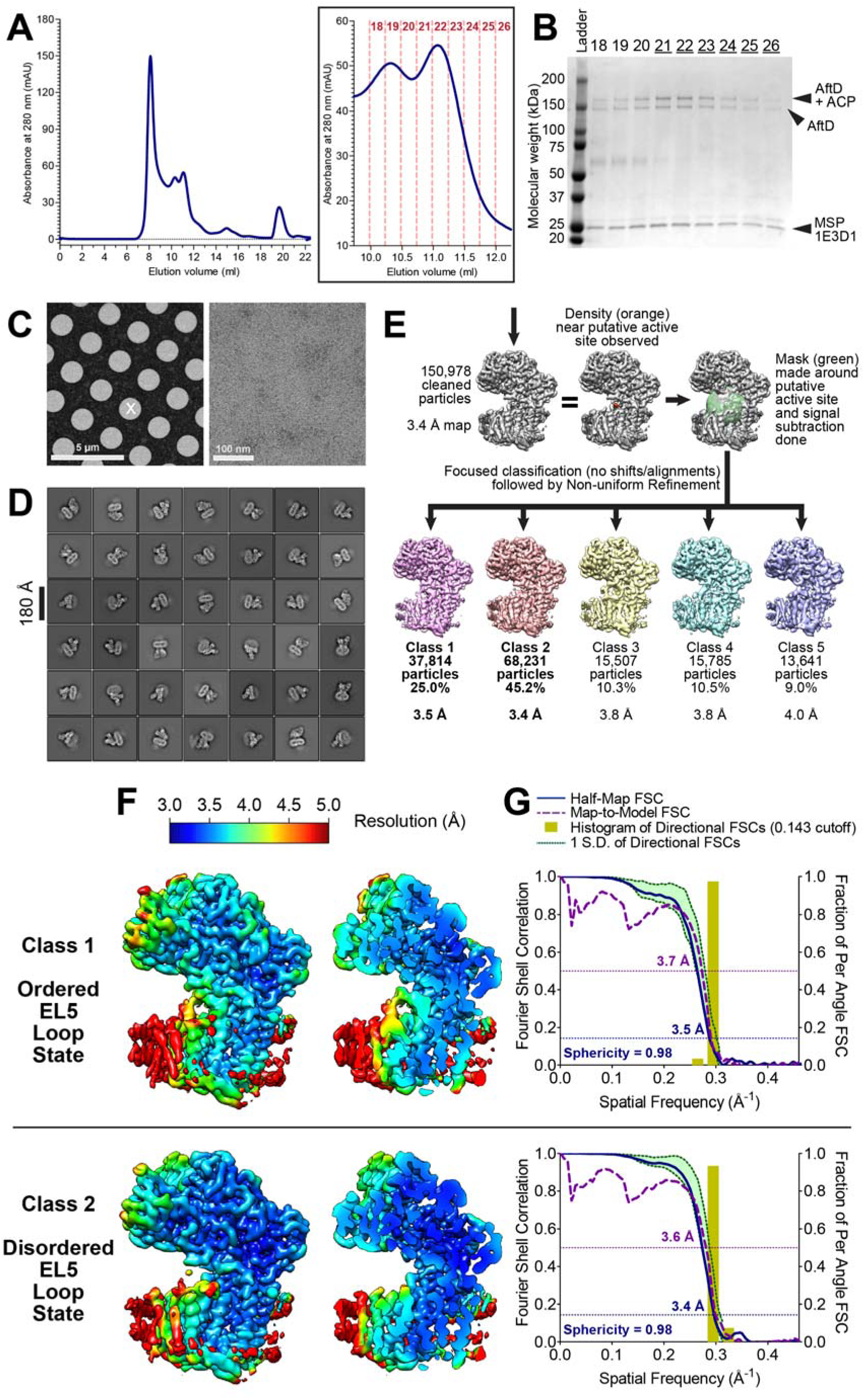
Purification and single-particle cryo-EM structural determination of AftD-R1389S, Related to Figure 6. (A) Representative size-exclusion chromatography (SEC) trace of AftD-R1389S incorporated into MSP-1E3D1 nanodiscs. The insert shows a zoom in of the fractions collected for SDS-PAGE analysis. (B) SDS-PAGE gel of the concentrated sample from the peak fractions stained with Coomassie blue. The predicted molecular weight of AftD-R1389S is 150 kDa, while that for MSP-1E3D1 is 32 kDa. Mass spectrometry was used to determine that the higher band at around 150 kDa corresponds to AftD-R1389S with ACP still associated. (C) Representative square, hole and exposure magnification views of the gold grids that were imaged. The exposure image had a defocus of -1.6 μm (estimated by CTFFind4) and pixel size of 1.0605 Å/pixel. (D) Representative 2D class averages. (E) Signal subtraction and 3D focused classification scheme. (F) ResMap display of two classes from focused classification of unsharpen AftD R1389S. (G) Fourier shell correlation (FSC) curves for the two classes describing the half-map (blue) resolutions (at 0.143 cut-offs), map-to-model (purple) resolutions (at 0.5 cut-offs), as well as histogram of directional resolutions sampled evenly over the 3DFSC (yellow). The corresponding sphericity value is also indicated.

**Table S1.**
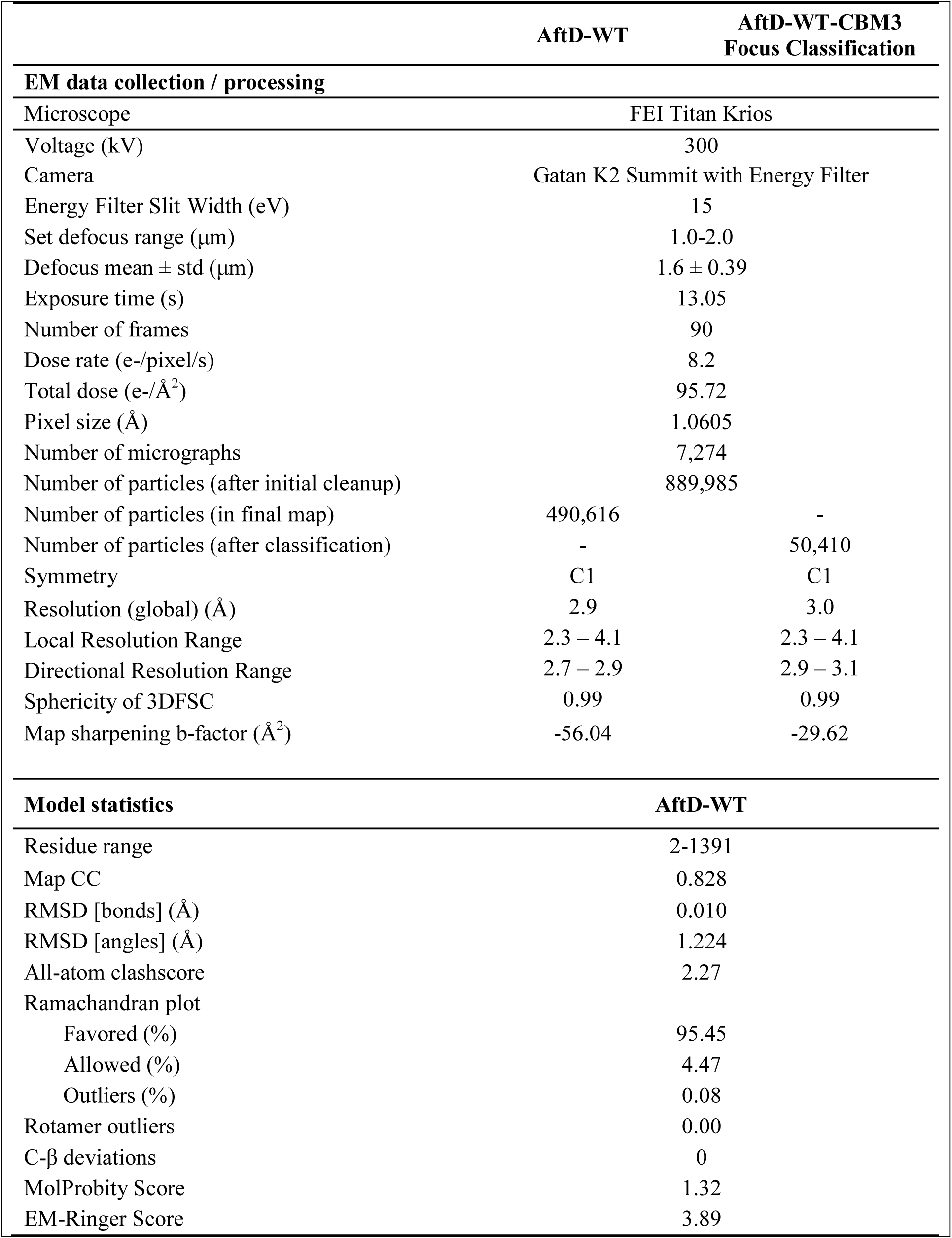
Cryo-EM data collection and modeling statistics WT-AftD, Related to Figure 2.

**Table S2.** Proteins identified by Liquid MS/MS from M. smegmatis purified AftD-WT M. abscessus, Related to Figure 5.

| Name | Accession UniProt | Unused ProtScore | Total ProtScore | Confidence (95%) | Species | Number of Peptides (95%) |
| --- | --- | --- | --- | --- | --- | --- |
| ATP synthase subunit beta | sp\|A0R200\|ATPB_MYCS2 | 62,02 | 62,02 | 66,1 | *M. smegmatis* | 31 |
| UPF0182 protein MSMEG_1959/MSMEI_ | sp\|A0QTT7\|Y1959_MYCS2 | 38,08 | 38,08 | 25 | *M. smegmatis* | 21 |
| ATP synthase subunit alpha | sp\|A0R202\|ATPA_MYCS2 | 37,14 | 37,19 | 50,4 | *M. smegmatis* | 19 |
| **AftD Mycobacterium abscessus 1948**<br>**F5/8 type C domain protein** |  | **32,81** | **32,81** | **11,4** | ***M. abscessus*** | **18** |
| 60 kDa chaperonin 1 | sp\|A0QQU5\|CH601_MYCS2 | 28,05 | 28,05 | 39,2 | *M. smegmatis* | 14 |
| Adenosylhomocysteinase | sp\|A4ZHR8\|SAHH_MYCS2 | 24,52 | 24,52 | 34,2 | *M. smegmatis* | 14 |
| Elongation factor Tu | sp\|A0QS98\|EFTU_MYCS2 | 24,42 | 24,45 | 48 | *M. smegmatis* | 13 |
| Probable malate:quinone oxidoreductase | sp\|A0QVL2\|MQO_MYCS2 | 18,94 | 18,94 | 24,7 | *M. smegmatis* | 10 |
| Cell wall synthesis protein Wag31 | sp\|A0R006\|WAG31_MYCS2 | 16,6 | 16,6 | 34,9 | *M. smegmatis* | 8 |
| ATP synthase gamma chain | sp\|A0R201\|ATPG_MYCS2 | 12 | 12 | 30 | *M. smegmatis* | 6 |
| Ketol-acid reductoisomerase (NADP(+)) | sp\|A0QUX8\|ILVC_MYCS2 | 11,91 | 11,95 | 22,9 | *M. smegmatis* | 6 |
| Glutamine synthetase | sp\|A0R079\|GLN1B_MYCS2 | 9,83 | 9,83 | 12,3 | *M. smegmatis* | 5 |
| Galactofuranosyltransferase GlfT2 | sp\|A0R628\|GLFT2_MYCS2 | 8,02 | 8,02 | 9,4 | *M. smegmatis* | 4 |
| 30S ribosomal protein S3 | sp\|A0QSD7\|RS3_MYCS2 | 8,01 | 8,01 | 15,3 | *M. smegmatis* | 4 |
| Uncharacterized protein<br>MSMEG_4692/MSMEI_4575 | sp\|A0R1B5\|Y4692_MYCS2 | 8 | 8 | 44 | *M. smegmatis* | 4 |
| ATP synthase epsilon chain | sp\|A0R1Z9\|ATPE_MYCS2 | 8 | 8 | 41,3 | *M. smegmatis* | 4 |
| 30S ribosomal protein S5 | sp\|A0QSG6\|RS5_MYCS2 | 6 | 6 | 25,7 | *M. smegmatis* | 3 |
| **Acyl carrier protein** | tr\|A0A0D6IL69\|A0A0D6IL69_MYCSM | **6** | **6** | **53,5** | ***M. smegmatis*** | **3** |
| Pyruvate dehydrogenase E1 component | sp\|A0R0B0\|ODP1_MYCS2 | 5,81 | 5,87 | 3,8 | *M. smegmatis* | 3 |
| Decaprenylphosphoryl-2-keto-beta-D-erythro-pentose reductase | sp\|A0R610\|DPRE2_MYCS2 | 5,07 | 5,13 | 12,6 | *M. smegmatis* | 3 |
| Acetamidase | sp\|Q07838\|AMDA_MYCSM | 4,01 | 4,02 | 5,4 | *M. smegmatis* | 2 |
| 50S ribosomal protein L5 | sp\|A0QSG1\|RL5_MYCS2 | 4 | 4 | 14,4 | *M. smegmatis* | 2 |
| ATP synthase subunit b | sp\|A0R204\|ATPF_MYCS2 | 2,29 | 2,29 | 7,6 | *M. smegmatis* | 1 |

**Table S3.** Cryo-EM data collection and modeling statistics AftD R1389S, Related to Figure 6.

|  | AftD R1389S<br>Class 1 | AftD R1389S<br>Class 2 |
| --- | --- | --- |
| **EM data collection / processing** |  |  |
| Microscope | FEI Titan Krios |  |
| Voltage (kV) | 300 |  |
| Camera | Gatan K2 Summit with Energy Filter |  |
| Energy Filter Slit Width (eV) | 15 |  |
| Set defocus range ($\mu\text{m}$) | 0.8-1.6 | |
| Defocus mean ± std ($\mu\text{m}$) | $1.42 \pm 0.35$ | |
| Exposure time (s) | 13.05 |  |
| Number of frames | 90 |  |
| Dose rate (e-/pixel/s) | 8.34 |  |
| Total dose (e-/$\text{\AA}^2$) | 96.78 | |
| Pixel size ($\text{\AA}$) | 1.0605 | |
| Number of micrographs | 4,886 |  |
| Number of particles (after initial cleanup) | 226,478 |  |
| Number of particles (in consensus map) | 150,978 |  |
| Number of particles (after classification) | 37,814 | 68,231 |
| Symmetry | C1 | C1 |
| Resolution (global) ($\text{\AA}$) | 3.5 | 3.4 |
| Local Resolution Range | 3.0 – 8.7 | 3.0 – 7.2 |
| Directional Resolution Range | 3.3 – 3.6 | 3.2 – 3.6 |
| Sphericity of 3DFSC | 0.98 | 0.98 |
| Map sharpening b-factor ($\text{\AA}^2$) | -125.09 | -129.19 |
| **Model statistics** |  |  |
| Residue range | 5-1388 | 5-1388 |
| Map CC | 0.8403 | 0.8045 |
| RMSD [bonds] ($\text{\AA}$) | 0.012 | 0.010 |
| RMSD [angles] ($\text{\AA}$) | 1.364 | 1.310 |
| All-atom clashscore | 4.12 | 3.23 |
| Ramachandran plot |  |  |
| Favored (%) | 92.12 | 93.62 |
| Allowed (%) | 7.80 | 6.30 |
| Outliers (%) | 0.08 | 0.08 |
| Rotamer outliers | 0.11 | 0.32 |
| C-β deviations | 0 | 0 |
| MolProbity Score | 1.68 | 1.54 |
| EM-Ringer Score | 3.70 | 3.21 |

## METHODS AND RESOURCES

### KEY RESOURCES TABLE

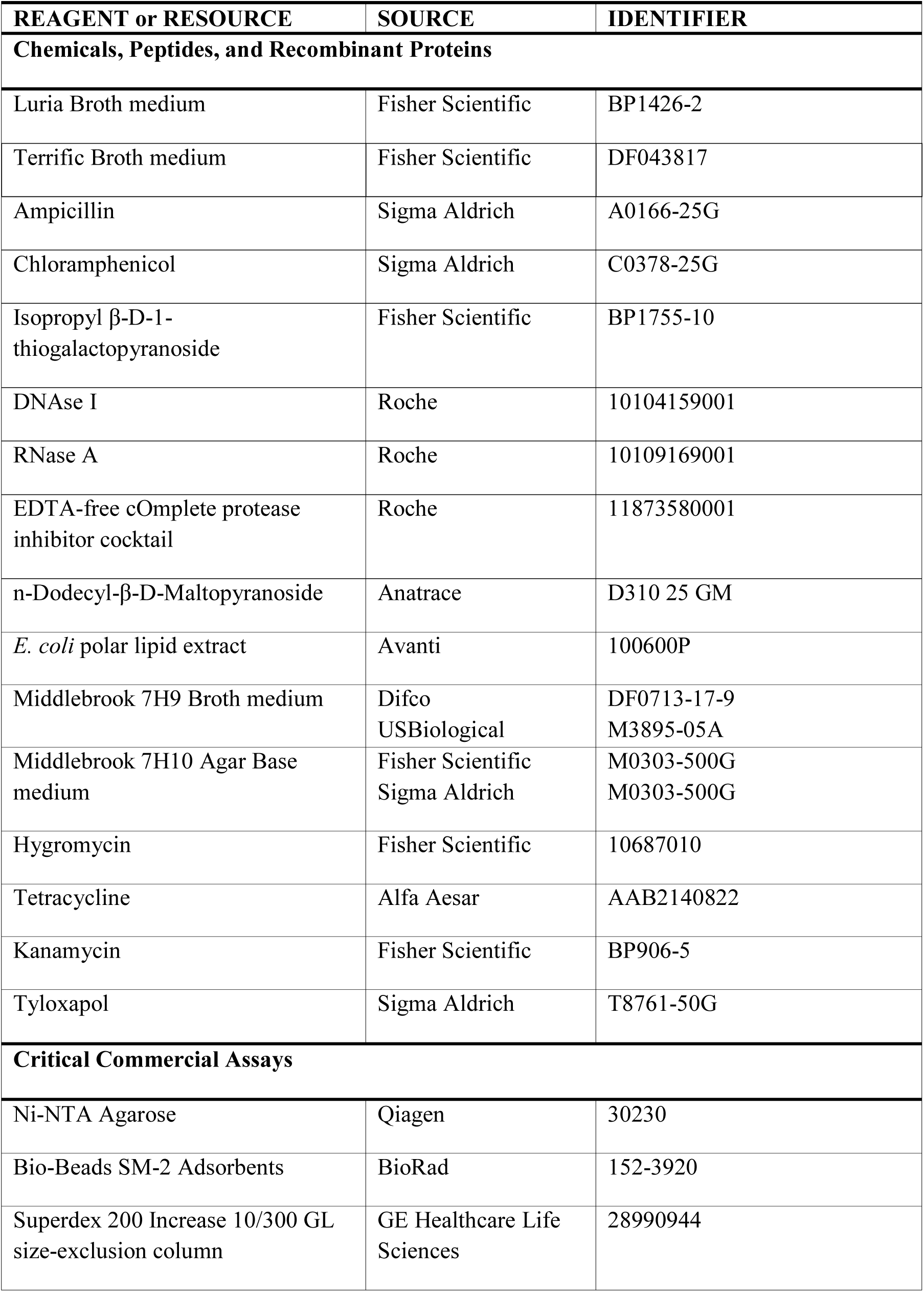
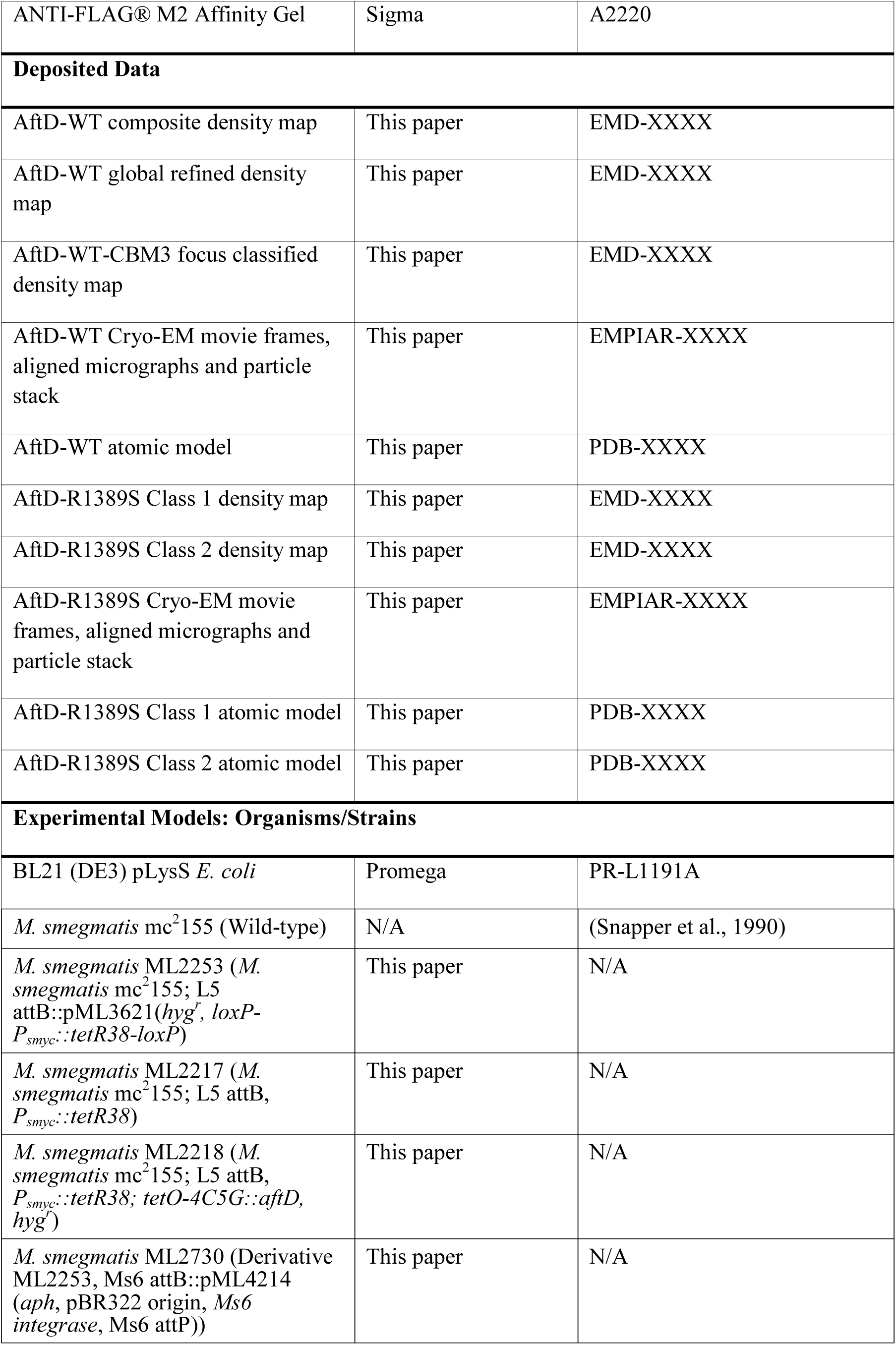
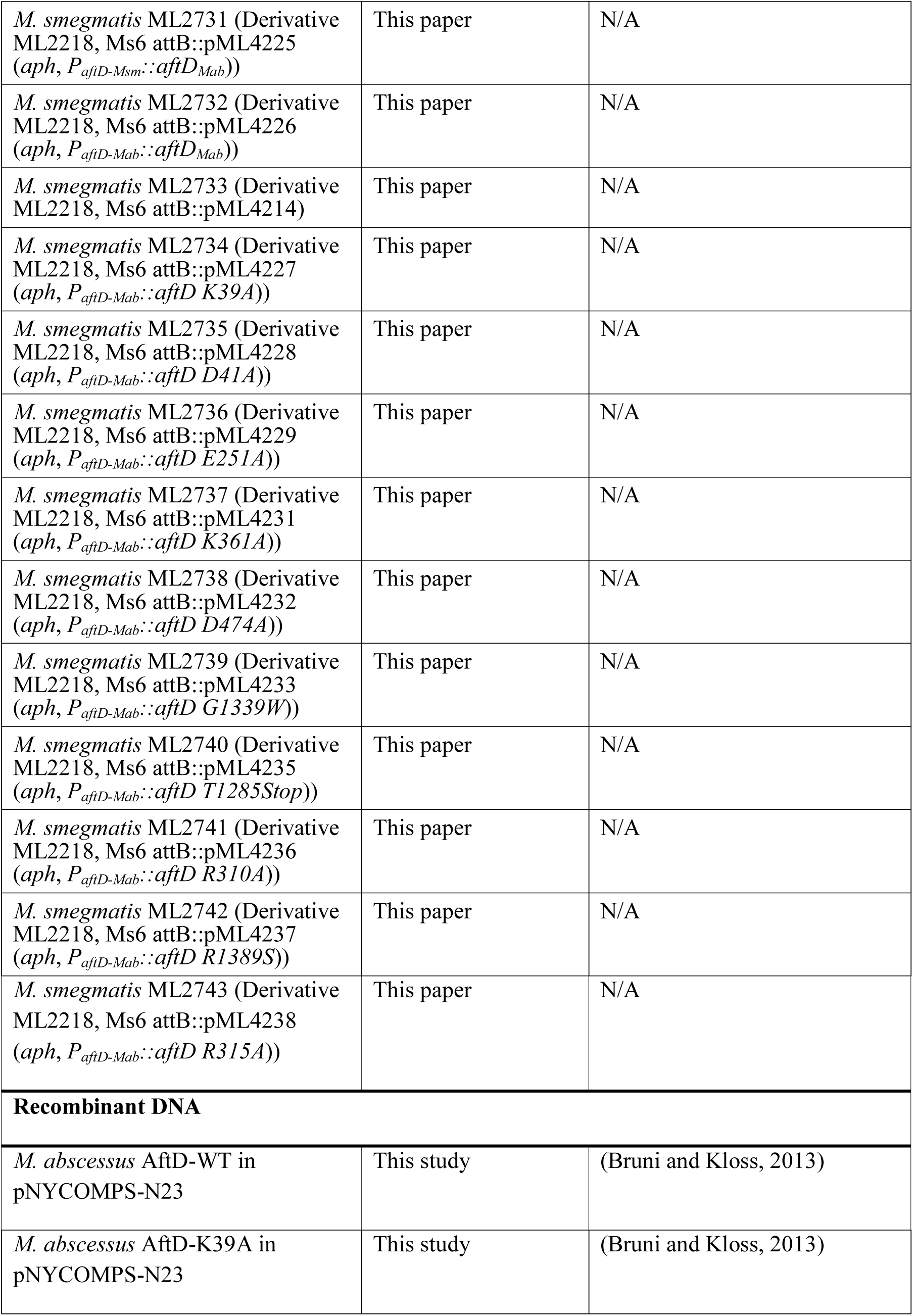
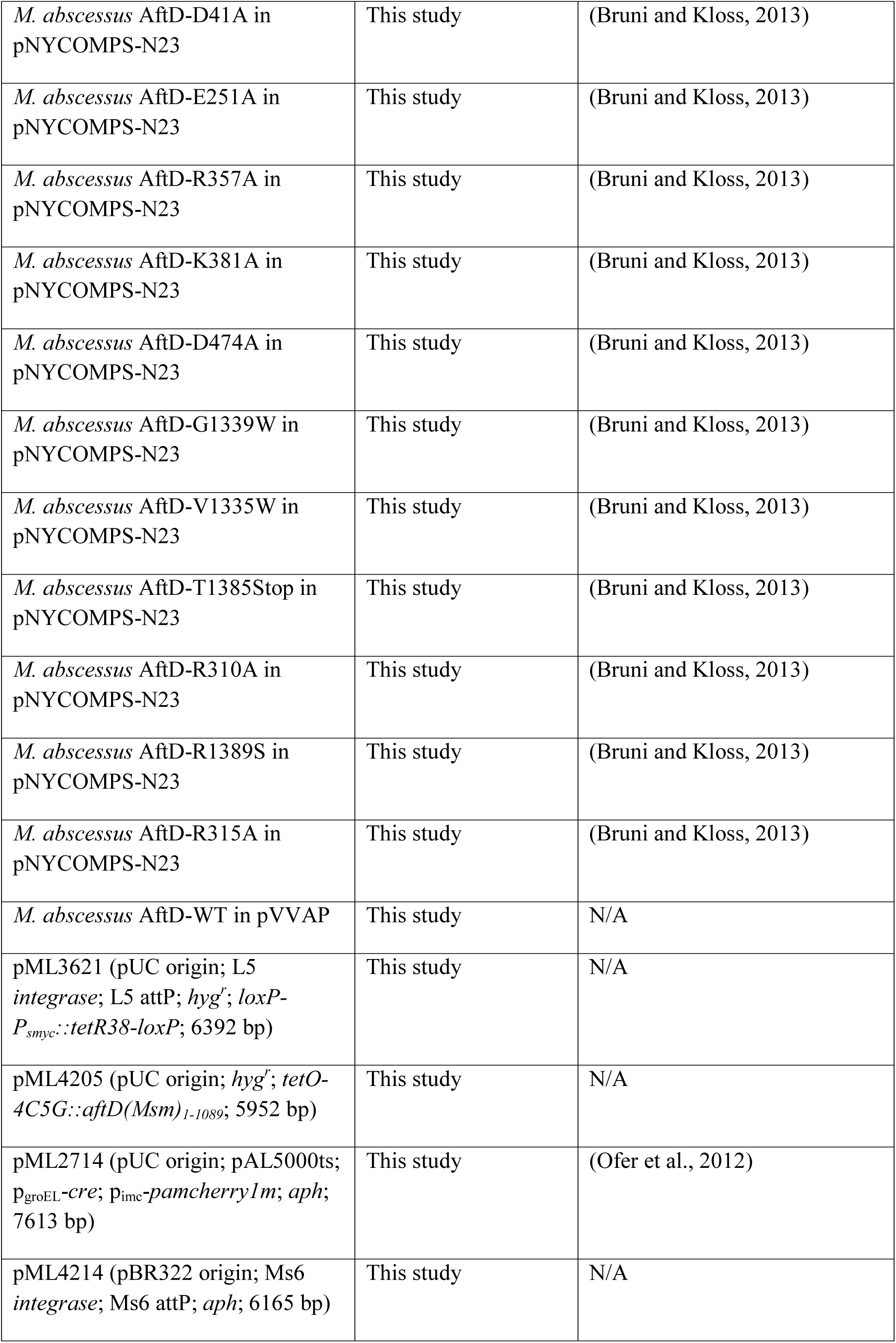
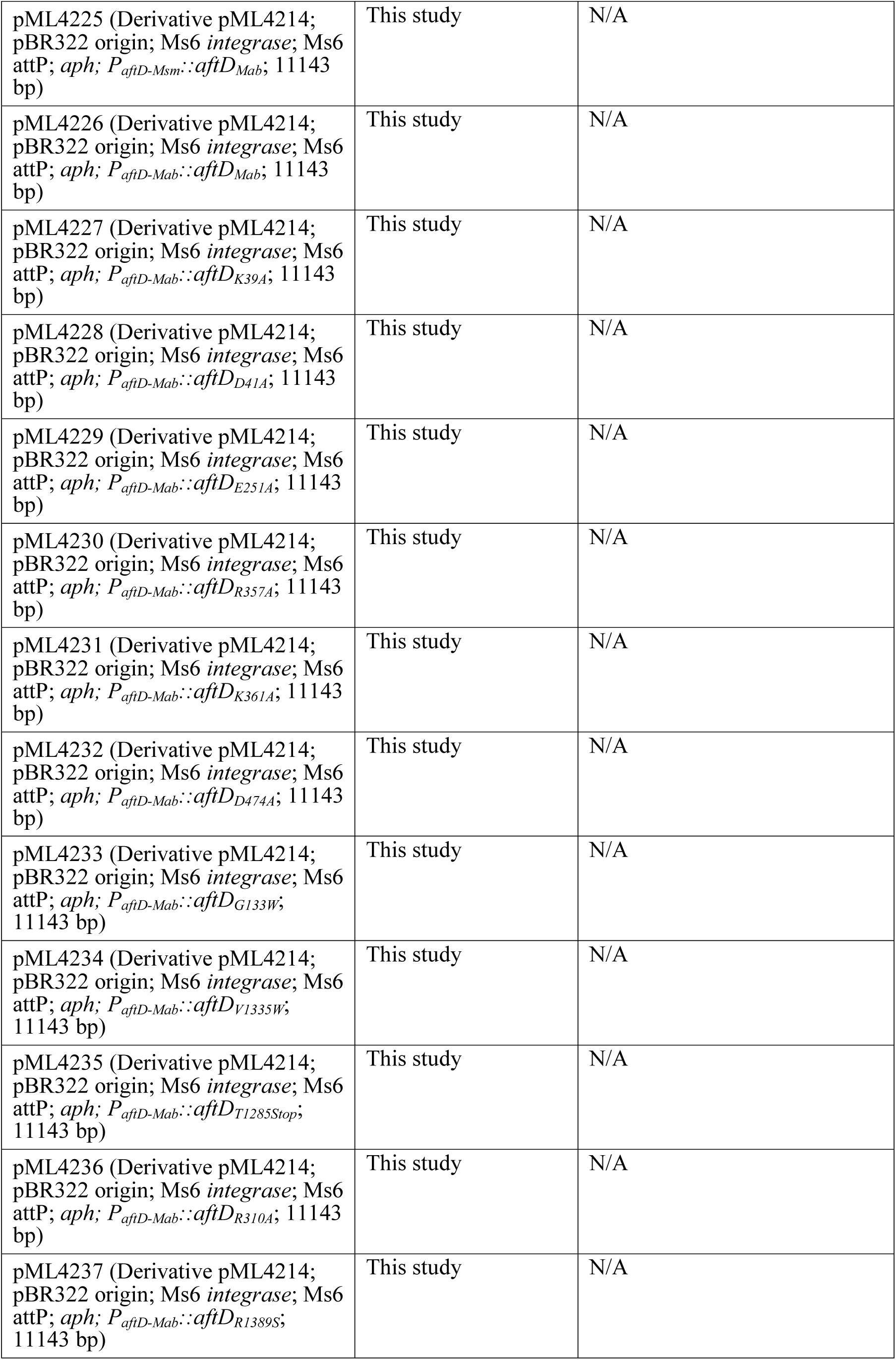
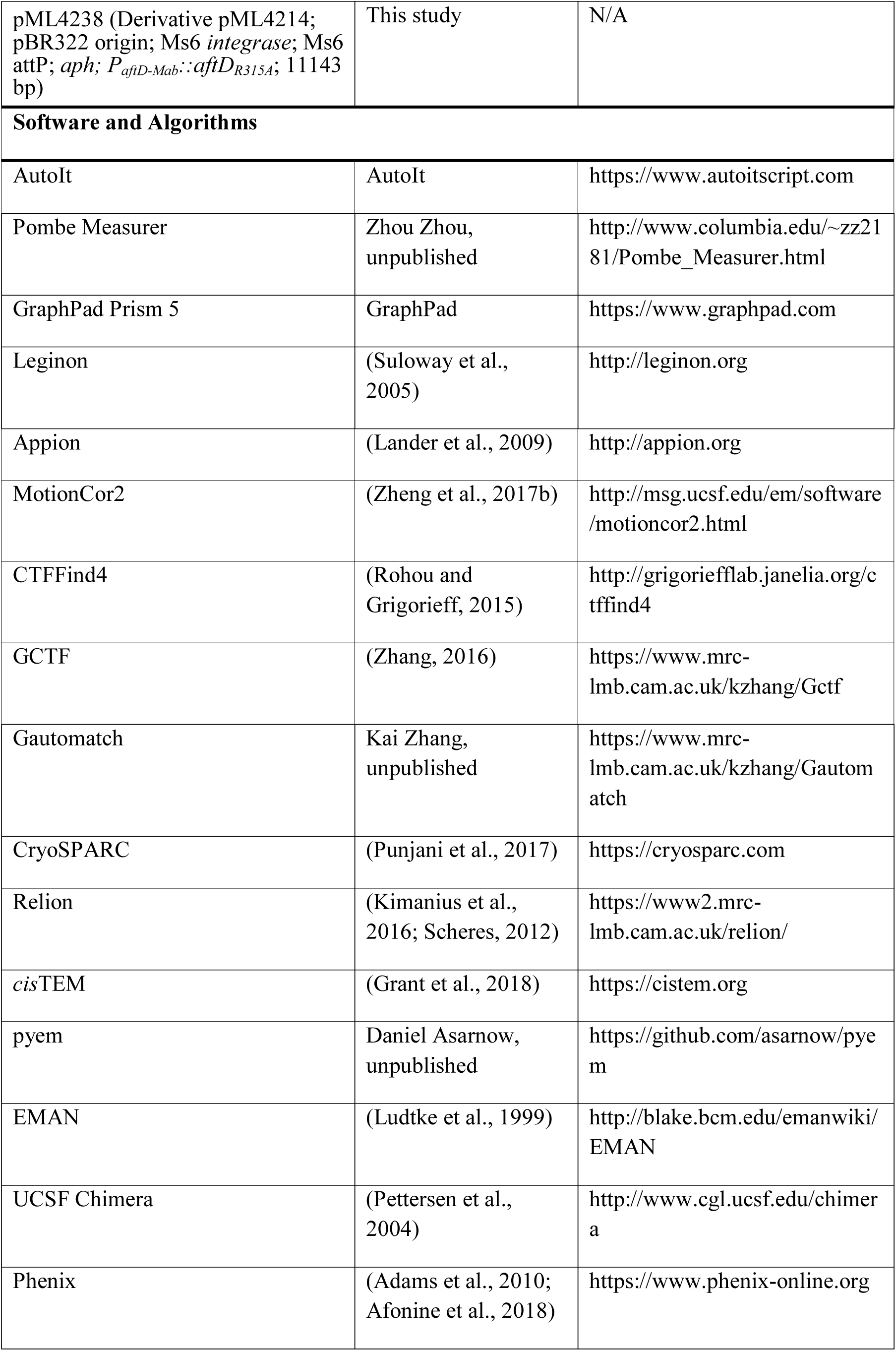
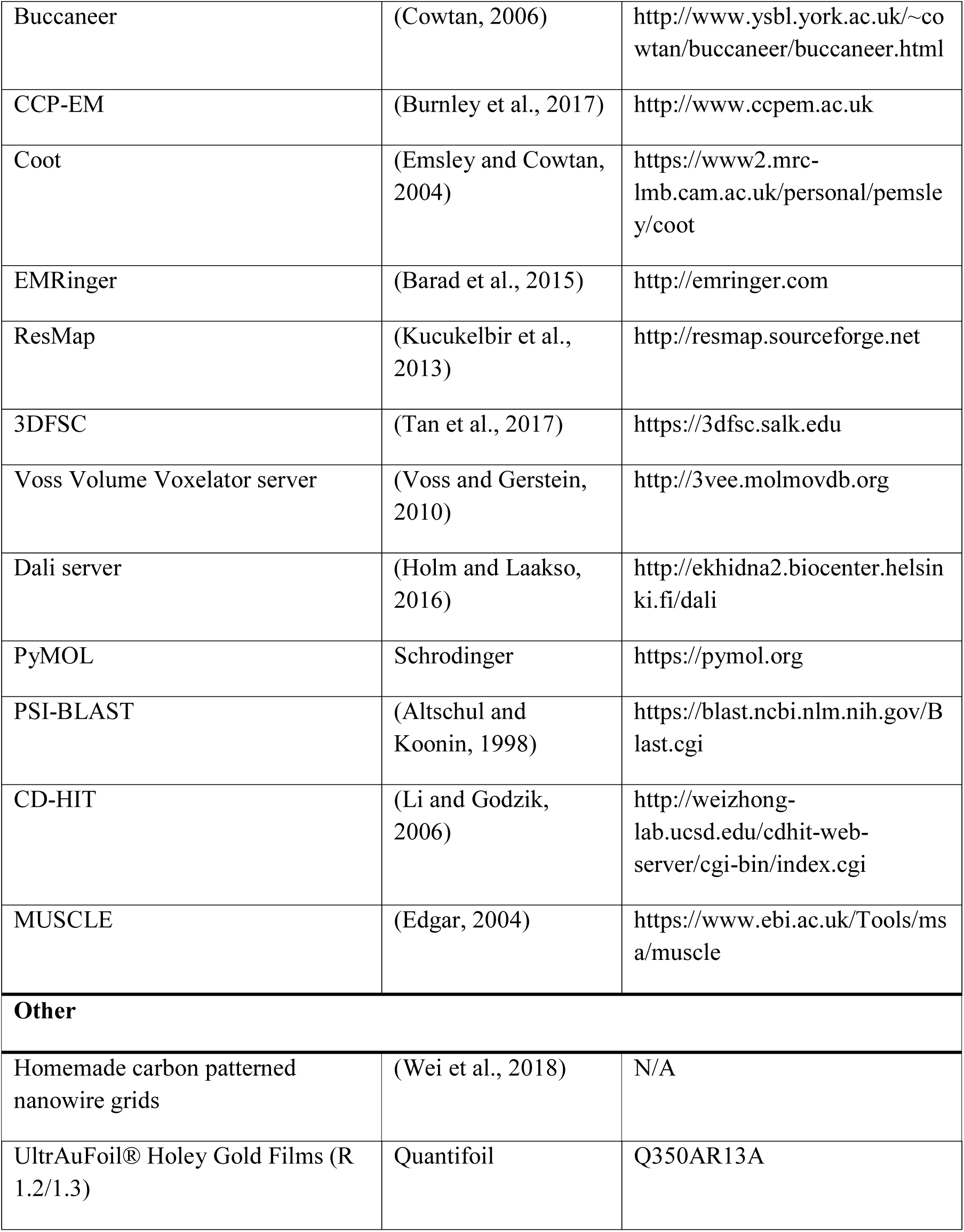

### CONTACT FOR REAGENT AND RESOURCE SHARING

Further information and requests for reagents may be directed to, and will be fulfilled by Dr. Filippo Mancia.

## Methods

### Statistics

For calculations of Fourier shell correlations (FSC), the FSC cut-off criterion of 0.143 (Rosenthal and Henderson, 2003) was used. No statistical methods were used to predetermine sample size. The experiments were not randomized. The investigators were not blinded to allocation during experiments and outcome assessment. For calculation of *M. smegmatis* cell length and width, both D’Agostino & Pearson omnibus normality test and Shapiro-Wilk normality test showed that the distribution was not normal. Hence, one-way ANOVA Kruskal-Wallis parametric test was done. Dunns post-test was then done between all pairs of samples.

### Phylogenetic and Conservation Analysis

PSI-BLAST (Altschul and Koonin, 1998) was performed using the amino acid sequence of *M. abscessus* AftD until no more additional sequences were found, producing 2385 sequences. Duplicate sequences from the same species and truncated variants of the same protein were removed, resulting in 665 unique sequences. Lengths of these sequences were plotted in Figure S1. Membrane topology for these sequences were predicted using the TMHMM server (Krogh et al., 2001).

In order to compare sequences that are more closely related to *Mycobacterium* AftD, sequences shorter than 95% the length of *M. abscessus* AftD (∼1400 residues) were excluded. CD-HIT (Li and Godzik, 2006) was then used to provide representative sequences and remove biases in the distribution of the data – notably *Mycobacterium* sequences dominate the original sequence list likely because of the medical relevance of this genus. The 289 final sequences were then aligned with MUSCLE (Edgar, 2004) and used for conservation analysis.

### Generation of a *M. smegmatis aftD* Conditional Knockout Strain

To construct a parent strain for *aftD* conditional knockout (cKO), a L5 integration vector pML3621 (L5 *integrase*, *Hyg^R^, loxP-P_smyc_::tetR38-loxP*) was firstly transformed into wild-type *M. smegmatis* mc^2^155 cell to create ML2253 strain which can constitutively express a reverse TetR (TetR38). The ML2253 colony was selected on 7H10 plates supplemented with 50 μ L hygromycin (Fisher) after 4 days at 37°C. Subsequently, a temperature-sensitive replication vector pML2714 (pAL5000ts, *Kan^R^*, *cre* recombinase expression) was transformed into ML2253 for unmarking. The unmarked colony (named ML2217) was selected on 7H10 plates supplemented with 30 μ mL kanamycin (Fisher) after 7 days at 32°C. Next, the unmarking plasmid ML2714 was removed by temperature selection on 7H10 (no antibiotic) plates at 40°C for 4 days.

To conditionally knockout *aftD* in *M. smegmatis* mc^2^155, a suicide vector pML4205 containing a TetR38-binding region (tetO-4C5G) and an N-terminus of *aftD* gene (1^st^ to 1089^th^ bp) was transformed into the parent strain ML2217 for single cross-over selection on 7H10 plates μg/mL hygromycin at 37°C for 4 days. PCR validation was then performed for the single cross-over colonies. The correct colonies were named ML2218, the *aftD* conditional KO strain. In the presence of 20 ng/mL tetracycline, TetR38-tetracycline complex binds the upstream region (tetO-4C5G) of *aftD* and represses the transcription of *aftD* gene.

### *M. abscessus aftD* Complementation

For the complementation experiment, the *M. abscessus aftD* gene, as well as the *aftD* variations, were cloned into an Ms6 integration vector pML4214 (Ms6 *integrase*, Ms6 attP attachment site, *Kan^R^*) individually with a 200-bp upstream DNA fragment of *M. abscessus aftD* gene. The pML4214 and the pML4214-derivative plasmids were transformed into ML2218 strain individually. All the colonies were selected on 7H10 plates supplemented with 30 μ /mL kanamycin (Fisher) after 4 days at 37°C.

### Phenotypic Assays for *M. smegmatis aftD* Conditional Knockout Strain

To observe a clear phenotype of *aftD* conditional knockout mutant, the remaining cellular AftD protein levels were depleted as much as possible. First, the ML2218 seed culture was grown to an OD_600_ of 1.0 at 37°C with a shaking speed of 200 rpm. For the AftD depletion, ML2218 cells were grown in 10 mL of 7H9/glycerol/tyloxapol liquid media supplemented with 20 ng/mL tetracycline with an initial OD_600_ of 0.002 at 37°C, 200 rpm for 2-3 days. Subsequently, the culture was used for growth curve assay, microplate appearance assay and colony forming assay.

For the growth curve assay, all the AftD-depleted strains were inoculated into the fresh 10 mL of 7H9/glycerol/tyloxapol liquid media supplemented with or without 20 ng/mL tetracycline with an initial OD_600_ of 0.005. The cultures were then grown at 37°C, 200 rpm and time points taken approximately every 4-8 hours till stationary phase. All experiments were done in triplicate.

For the microplate appearance assay, all the AftD-depleted strains were inoculated into 12-well plates (Costar). Each well contains 3 mL of 7H9/glycerol/tyloxapol liquid media supplemented with or without 20 ng/mL tetracycline with an initial OD_600_ of 0.05. The 12-well plates were shaking at 120 rpm at 37°C for 2-3 days.

For the colony forming assay, all the AftD-depleted cells were streaked on 7H10 plates with or without 20 ng/mL of tetracycline. The plates were kept at 37°C for 6 days.

### Scanning Electron Microscopy

10mL of *M. smegmatis* cells are harvested by spinning cells at 8000g for 1 min with a Sorvall ST 16 R centrifuge (Thermo Scientific) to give around 0.4-0.5g of cell pellet mass. Supernatant was discarded. Cell pellet was then resuspended with 2.5% glutaraldehyde (Tousimis) in a 15mL Falcon tube and incubated at 4°C for 2 hours. The suspension was then spun down again at 8000g for 1 min and resuspended with PBS for 2 times, then with an ethanol (Fisher) gradient (30%, 50%, 70%, 100%) 15 min for dehydration. Each step takes 15 minutes, and will be followed with a spin down and resuspension. After the final dehydration, fixed cells are left to air dry overnight.

The cells were then smeared onto a scanning electron microscope sample holder using a pipette tip for imaging. The entire surface of the specimen was then sputter coated with a thin layer of gold/palladium. The cells were imaged using secondary electron mode in a FEI Helios Nanolab 650 (FEI). Images were recorded using the scanning electron microscope beam at 2 keV and 100 pA with a working distance of 4.0 ± 0.5 mm. Data acquisition occurred manually using an in-house AutoIt script (https://www.autoitscript.com/) written by Bill Rice. The raw images were 3072 pixels by 2048 pixels, with 6.08 nm pixel width, a horizontal field width (HFW) of 18.67 µm and dwell time of 10.00 µs.

To measure the dimensions of cells, Adobe Photoshop was used to select out each cell. The cell length and width were calculated using ImageJ’s plugin, Pombe Measurer (Zhou Zhou, www.columbia.edu/~zz2181/Pombe_Measurer.html). Both D’Agostino & Pearson omnibus normality test and Shapiro-Wilk normality test showed that the distribution of lengths and widths were not normal. Hence, one-way ANOVA Kruskal-Wallis parametric test was done. Dunns post-test was then done between all pairs of samples. Statistical tests were done using GraphPad Prism 5.

### Genomic Expansion and Small-Scale Screening

AftD genes were identified from a collection of fourteen *Mycobacterium* genomes using a bioinformatics approach (Love et al., 2010). Ligation independent cloning (LIC) was used to clone these targets from the genomes into five LIC-adapted expression vectors (pNYCOMPS-Nterm, pNYCOMPS-Cterm, pNYCOMPS-N23, pNYCOMPS-C23 and pMCSG7-10x) that contained a Tobacco Etch Virus (TEV) protease cleave site (ENLYFQSYV) and decahistidine affinity tag. Small and medium scale expression was done in a high throughput manner as described in detail in a previous protocol by Bruni & Kloss (Bruni and Kloss, 2013). Only one construct, *M. abscessus* AftD in the pNYCOMPS-N23 plasmid, was successfully cloned and expressed in small and medium scale. This construct was thus brought forward for further structural studies.

### *M. abscessus* AftD Mutagenesis

Site-directed mutagenesis on the *M. abscessus aftD* gene was done with overlapping primers designed per mutant using the Quick Change Lightning Kit (Agilent). The same pairs of primers were used on *aftD* in pNYCOMPS-N23 plasmid (for *E. coli* expression) and pML4214-Mab Promoter::*aftD M. abscessus* (for *M. smegmatis* expression).

Small scale expression tests were carried out for the AftD mutants in the pNYCOMPS-N23 plasmid as described in detail in a previous protocol by Bruni & Kloss (Bruni and Kloss, 2013) (Figure S6). No TEV digestion was carried out for the small scale expression tests.

### *M. abscessus* AftD expression, purification and nanodisc reconstitution

*M. abscessus* AftD WT in the pNYCOMPS-N23 plasmid was transformed into BL21 (DE3) pLysS *E. coli* competent cells and plated onto Luria broth (LB) agar (Fisher) plates supplemented with 100 μg/mL chloramphenicol (Sigma) and grown overnight at 37°C. In the next day, a colony was picked and used to inoculate a starter culture containing 150 mL of Terrific broth (TB) medium (Fisher) supplemented with 100 μg/mL chloramphenicol. The starter culture was grown overnight at 37°C with shaking (240 r.p.m.) in an incubator shaker (New Brunswick Scientific). In next day, 12 2L baffled flasks each with 800 mL of Terrific broth (TB) medium (Fisher) supplemented with 100 μ g/mL chloramphenicol were inoculated with 10 mL of starter culture. The cultures were then grown at 37°C with shaking (240 r.p.m.) until optical density (OD) at 600nm reached 1.0 (approximately 3 hours). Temperature was then reduced to 22°C and AftD expression was induced with 0.2 mM β-D-1-thiogalactopyranoside (IPTG) (Fisher). The culture was then incubated overnight with shaking (240 r.p.m.). In the next day, the cells were harvested by centrifugation at 4,000 g in H6000A/HBB6 rotor (Sorvall) for 30 min at 4°C. The supernatant was discarded and the pellet was resuspended in chilled 1x phosphate buffered saline (PBS) and centrifuged again at 4,000 g for 30 min at 4°C. The supernatant was again discarded and the pellet was resuspended in lysis buffer containing 20 mM HEPES pH 7.5, 200 mM NaCl, 20 mM MgSO_4_, 10 μg/mL DNase I (Roche), 8 μg/mL RNase A (Roche), 1 mM tris(2-carboxyethyl)phosphine hydrochloride (TCEP), 1 mM PMSF, 1 tablet/1.5 L buffer EDTA-free cOmplete protease inhibitor cocktail (Roche). For 9.6 L of culture, there was ∼15-20g of wet cell pellet mass, which was resuspended with ∼250 mL of lysis buffer. Cells were lysed by putting the suspension through a chilled Emulsiflex C3 homogenizer (Avestin) three times. To isolate the membrane fractions, the lysate was put through ultracentrifugation at 37,000g in Type 45 Ti Rotor (Beckman Coulter) at 4°C for 30 min. The supernatant was discarded and the pellet was resuspended in the lysis buffer up to a volume of 240 mL and homogenized using a hand-held glass homogenizer (Konte). The membrane fraction was then stored at −80°C until the use.

The thawed membrane fraction was solubilized by adding n-dodecyl-β-D-maltopyranoside (DDM) to a final concentration of 1% (w/v) detergent for 2 hours at 4°C with gentle rotation. Insoluble material was removed by ultracentrifugation at 40,000g in Type 45 Ti Rotor at 4°C for 30 min. The supernatant was added to six Falcon tubes containing pre-equilibrated Ni^2+^-NTA resin (Qiagen) in the presence of 40 mM imidazole and incubated with gentle rotation at 4°C for 2 hours. The resin was washed with 10 column volumes of wash buffer containing 20mM HEPES pH 7.5, 200 mM NaCl, 60mM Imidazole, 0.1% DDM and eluted with elution buffer containing 20 mM HEPES pH 7.5, 200mM NaCl, 200mM Imidazole, 0.05% DDM. The eluted protein was exchanged into a buffer containing 20 mM HEPES pH 7.5, 200mM Imidazole, 0.05% DDM using a PD-10 desalting column (GE), and concentrated down using a 100-kDa concentrator (Pierce) to ∼ 1 mg/mL.

The protein was then incorporated into lipid nanodisc (Bayburt and Sligar, 2010) with the molar ratio 1:360:6 of *E. coli* polar lipid extract (Avanti): membrane scaffold protein 1E3D1 (MSP1E3D1) and incubated for 2 hours with gentle agitation at 4°C. Reconstitution was initiated by removing detergent with the addition of 150 mg Bio-beads (Bio-Rad) per mL of protein solution for overnight with constant rotation at 4°C. Bio-beads were removed by passing the protein solution through an Ultrafree centrifugal filter unit (Fisher) and the nanodisc reconstitution mixture was re-bound to Ni^2+^-NTA resin with 15 mM imidazole for 2 hours at 4°C in order to remove free nanodisc. The resin was washed with 10 column volumes of wash buffer consisting of 20 mM HEPES pH 7.5, 200 mM NaCl and 20 mM Imidazole, followed by 3 column volumes of elution buffer consisting of 20 mM HEPES pH 7.5, 200 mM NaCl and 200 mM Imidazole.

TEV protease (Kapust et al., 2002) (∼0.5 mg TEV protease added per pellet equivalent from 800 mL of initial bacterial culture) was added to the eluted protein, which was then injected into a Slide-A-Lyzer™ dialysis cassette (Thermo Fisher) and dialyzed against 200 mL of 20 mM HEPES pH 7.5 and 200 mM NaCl overnight at 4°C. The mixture was then re-bound to Ni^2+^-NTA resin with 15 mM imidazole for 1 hour at 4°C in order to remove uncleaved AftD, cleaved tag, TEV protease and contaminant proteins. The resin was washed with 3 column volumes of wash buffer consisting of 20 mM HEPES pH 7.5, 200 mM NaCl. The flowthrough and wash were concentrated using a 100-kDa concentrator to under 500 μL and loaded onto a Superdex 200 Increase 10/300 GL size-exclusion column (GE Healthcare Life Sciences) in gel filtration buffer (20 mM HEPES pH 7.5 and 200 mM NaCl). Throughout the entire process of purification, 15 μL of samples were taken and added to 5 μL of 6X reducing Laemmli SDS sample buffer (Bioland Scientific). The samples were then loaded on a 4–20% Mini-PROTEAN TGX precast protein gel (Bio-rad) for protein gel electrophoresis in a Tris/Glycine/SDS buffer. The gel was developed using InstantBlue (Sigma) protein stain.

For *M. abscessus* AftD-R1389S, the expression and purification protocol is largely the same except for the following. Firstly, AftD-R1389S was not able to be cleaved by TEV protease, hence the protein was not put through overnight TEV cleavage. Secondly, during the second nickel affinity purification after reconstitution into nanodiscs, additional washes of 5 column volumes of 20 mM HEPES pH 7.5, 500 mM NaCl and 20 mM Imidazole and 5 column volumes of 20 mM HEPES pH 7.5, 500 mM NaCl, 5 mM ATP and 20 mM Imidazole were done in order to increase purity of the protein, given TEV cleavage was no longer done.

### Single-Particle Cryo-EM Sample Vitrification

Purified AftD WT was concentrated using a 100-kDa concentrator (Pierce) to between 5-20 μ of sample at ∼1 mg/mL. Due to the low yields obtained, the samples were vitrified using a semi-automated Spotiton V1.0 robot (Dandey et al., 2018; Jain et al., 2012; Razinkov et al., 2016). This is a device for vitrifying cryo-EM samples that uses piezo-electric dispensing to apply small (50 pl) drops of sample across a ‘self-blotting’ nanowire grid as it flies past en-route to plunging into liquid ethane. Owing to the small amounts of sample required per vitrified grid, only ∼3 μL of sample was required to be aspirated for multiple grids. Homemade carbon support nanowire grids with a regular array of holes were used (Wei et al., 2018), and were plasma-cleaned (Gatan Solarus) for 15s using an H_2_/O_2_ mixture. The Spotiton V1.0 robot was operating at room temperature and moderate humidity.

For purified AftD-R1389S, the protein was concentrated using a 100-kDa concentrator (Pierce) to between ∼10 μL of sample was added to a plasma-cleaned (Gatan Solarus) 1.2/1.3 µm holey gold grid (Quantifoil UltrAuFoil) and blotted using filter paper on one side for 2s using the Leica GP plunger system before plunging immediately into liquid ethane for vitrification. The plunger was operating at 6°C with >80% humidity to minimize evaporation and sample degradation. Due to conditions previously optimized using Spotiton, Leica GP plunging could be used to obtain good grids for the mutant without expending too much of the protein sample.

### Data Acquisition

Images were recorded on a Titan Krios electron microscope (FEI) equipped with a K2 summit direct detector with a Quantum energy filter (Gatan) operating at 1.0605 Å per pixel in counting mode using the Leginon software package (Suloway et al., 2005). Pixel size was calibrated in-house using a proteasome test sample. Energy filter slit width of 15 eV was used during the collection and aligned automatically every hour using Leginon. Data collection was performed using a dose of ∼95.72 e^-^/Å^2^ across 90 frames (150 msec per frame) at a dose rate of ∼8.2 e^−^/pix/sec, using a set defocus range of −1.0 μ to −2.0 μ. 100 μm objective aperture was used. For AftD WT, a total of 7,274 micrographs were recorded over a four sessions using an image beam shift data collection strategy (Cheng et al., 2018). For AftD-R1389S, the same strategy over a single session of three days yielded 4,886 micrographs. The dose was marginally different at 96.78 e^−^/Å^2^.

### Data Processing

For *M. abscessus* WT AftD, data from the four sessions were processed separately and combined towards the end of the processing pipeline. Movie frames were aligned using MotionCor2 (Zheng et al., 2017b) with 5 by 5 patches and B-factor of 100 through the Appion software package (Lander et al., 2009). Micrograph CTF estimation was performed using both CTFFind4 (Rohou and Grigorieff, 2015) and GCTF (Zhang, 2016), and best estimate based on confidence was selected within the Appion software package. Template free particle picking with Gautomatch (Kai Zhang, unpublished, https://www.mrc-lmb.cam.ac.uk/kzhang/Gautomatch/) using an extremely lenient threshold (to avoid missing any particles) was used to pick 2,800,102 particles (extracted 256 box size binned by 2) that were transferred into Relion 2.1 (Kimanius et al., 2016; Scheres, 2012) for 2D classification. 2D class averages that were ice or showed no features were discarded. An initial model was then generated in CryoSPARC *ab initio* (Punjani et al., 2017) and used for Relion 3D refinement. The Euler angles and shifts were then used to re-extract the particles and re-center them. Relion 3D classification was then performed using 7 classes and only classes with clear structural features were selected.

At this stage, a total of 498,060 selected particles from all four session were combined and re-extracted unbinned. Per-particle CTF using GCTF was then performed before the particles were put through a Relion 3D refinement using the previously obtained initial model. A mask generated around the transmembrane region was then used to performed 3D classification with 8 classes, and particles belonging to classes with high resolution features were selected and re-extracted with a bigger box size of 288. A total of 490,616 particles were left in the final stack.

CryoSPARC non-uniform refinement produced a 3.2 Å map, which was then put through *cis*TEM (Grant et al., 2018) CTF refinement to obtain better defocus values. The particles with the refined defocus values were then put through another round of CryoSPARC non-uniform refinement to produce a 3.0 Å map. Signal subtraction was then done in Relion to remove the nanodisc and flexible ACP density, and the signal subtracted particles were then put through CryoSPARC non-uniform refinement. The resulting Euler angles and shifts were then used to reconstruct the non-signal subtracted particles to produce a 2.9 Å map, called the AftD global refined map. This final map was sharpened using phenix.auto_sharpen (Adams et al., 2010; Afonine et al., 2018), which automatically determined that b_iso sharpening to high_resolution cutoff should be the algorithm to use. Overall b_sharpen applied was 56.04 Å^2^ and final b_iso obtained was 44.30 Å^2^.

When examining the map by eye, it was noticed the domain corresponding to CBM3 had a lower resolution (poor density) compared to the rest of the map. In order to better improve the resolution at this flexible portion, signal subtraction was performed to isolate just the CBM3 region, and focused classification using ten classes without alignment was done. A total of 50,410 particles were present in the highest resolution class. The resulting Euler angles and shifts were then used to reconstruct the non-signal subtracted particles to produce a 3.0 Å map, called the AftD-CBM3 focused classified map. This final map was sharpened using phenix.auto_sharpen, which automatically determined that b_iso sharpening to high_resolution cutoff should be the algorithm to use. Overall b_sharpen applied was 29.62 Å^2^ and final b_iso obtained was 45.97 Å^2^.

For *M. abscessus* AftD-R1389S, the same data processing strategy was used except for the following differences. Firstly, 226,478 particles were picked initially and a consensus map of 150,978 particles was obtained. Secondly, from these 150,978 particles, signal subtraction was done using a spherical mask around the putative active site of AftD where additional density could be seen. Focused 3D classification without alignments was then performed using Relion with five classes. Different T values (10, 20, 30, 40, 50, 60, 70, 80 and 90) were titrated and T=60 was determined by visual inspection and comparison of classes to give the best classification. Classes 3, 4 and 5 looked noisy and not significantly different. Classes 1 and 2 had high resolution details and had significant differences between them. A CryoSPARC non-uniform refinement was then performed using particles from each of these classes to give final maps at 3.5 Å and 3.4 Å each respectively. Both final maps were sharpened using phenix.auto_sharpen, using b-factors of −125.09 Å^2^ and −129.19 Å^2^ respectively.

All conversions between Relion, CryoSPARC, and *cis*TEM were performed using Daniel Asarnow’s pyem script (unpublished, https://github.com/asarnow/pyem).

### Model Building and Refinement

In order to aid in model building against a single structure, phenix.combine_focused_maps (Adams et al., 2010; Afonine et al., 2018) was used to combine the WT AftD global refined map and WT AftD-CBM3 focused classified map into the WT AftD composite map.

For model building of WT AftD, the Chimera (Pettersen et al., 2004) program was first used to segment out the protein components and exclude the nanodisc. *De novo* model building for the cryo-EM map its primary sequence was initiated using Buccaneer (Cowtan, 2006) within the CCP-EM software suite (Burnley et al., 2017). After 2 rounds of Buccaneer model building, the partial model was brought into Coot (Emsley and Cowtan, 2004) program for manual model building. After the model was built, it was refined against the cryo-EM map utilizing real space refinement in the Phenix program. Restraints for the lipids were generated using phenix.eLBOW and for the metal ions using phenix.ready_set. Thereafter, model adjustment and refinement were performed iteratively in Coot and Phenix, with the statistics being examined using Molprobity (Chen et al., 2010) until no further improvements were observed. The final map and model were then validated using 1) EMRinger (Barad et al., 2015) to compare map to model, 2) ResMap (Kucukelbir et al., 2013) to calculate map local resolution and 3) 3DFSC program suite (Tan et al., 2017) to calculate degree of directional resolution anisotropy through the 3DFSC. Map-to-model FSCs were also calculated by first converting the model to a map using Chimera molmap function at Nyquist resolution (2.121 Å). A mask was made from this map using Relion (after low-pass filtering to 8 Å, extending by 1 pixel and applying a cosine-edge of 3 pixels), and was then applied to the density map. Map-to-model FSC was calculated using EMAN (Ludtke et al., 1999) proc3d between these maps.

For model building of AftD-R1389S, the WT structure was overlaid and Coot was used to manually remove or build residues. Other than that, the model building and refinement strategy mirrored the WT AftD mentioned above.

### Model Analysis

A cavity search using the Solvent Extractor from Voss Volume Voxelator server (Voss and Gerstein, 2010) was performed using an outer probe radius of 5 Å and inner probe radius of 2 Å. In order to search for other PDB structures with similar fold, a Dali server (Holm and Laakso, 2016) search was performed – first globally and then against the different domains of the model.

### Screening, Imaging and Data Analysis of the Glycan Array

Glycan array analysis was done with *M. abscessus* AftD WT protein solubilized in DDM. Slides were prewetted in buffer A (25 mM Tris-HCl pH 7.8, 0.15 mM NaCl, 2 mM CaCl_2_ and 0.05% Tween 20) for 5 min, rinsed with buffer B (25 mM Tris-HCl pH 7.8, 0.15 mM NaCl, and 2 mM CaCl_2_) three times, and blocked overnight with buffer C (1% BSA in 25 mM Tris-HCl pH 7.8, 0.15mM NaCl, and 2 mM CaCl_2_) at 4 °C. Aliquots (500 μL) of serial dilutions of protein samples in buffer C were transferred to wells of the slide module immediately after aspiration of the blocking buffer. Wells were sealed with an adhesive seal and incubated for 60 min at 37 °C. Protein was removed by aspiration, and slides were washed 10 times with buffer A and three times with buffer B. Fluorescence was measured directly or after addition of a secondary antibody in buffer C (1:1000 dilution). Slides were incubated with a secondary antibody at room temperature for 40 min before being washed repeatedly with buffer A and deionized water.

Before being scanned, slides were dried by centrifugation. Microarrays were scanned at 5 μ resolution with a GenePix 4000B scanner (Molecular Devices, Sunnyvale, CA). The fluorescent signal was detected at 532 nm for Cy3 or Alexa Fluor 555 and 488 nm for Alexa Fluor 488. The laser power was 100%, and the photomultiplier tube gain was 400. The fluorescent signals were analyzed by quantifying the pixel density (intensity) of each spot using GenePix ProMicroarray Image Analysis Software version 6.1. Fluorescence intensity values for each spot and its background were calculated. The local background signal was automatically subtracted from the signal of each separate spot, and the mean signal intensity of each spot was used for data analysis. Averages of triplicate experiments and standard deviations were calculated using Microsoft Excel.

### *M. smegmatis* Overexpression of M. abscessus AftD

*M. abscessus* AftD, containing N-terminal FLAG and His_10_ and tags, was cloned into pVVAP (Catalão et al., 2010) plasmid for over-expression. This plasmid contains a pAL5000 replication origin, a kanamycin resistance cassette, and allows expression of N terminally FLAG+His_10_ tagged AftD. The plasmid was transformed by electroporation into *M. smegmatis* MC^2^155 and plated onto Middlebrook 7H10 (Sigma-Aldrich), plates, supplemented with 0.5% glucose and 15 µg mL^-1^ kanamycin, and left at 37°C until single colonies were visible (2 to 3 days). A single colony was picked to inoculate 30 mL of Middlebrook 7H9 medium (USBiological) supplemented with 0.5% glucose, 0.05% Tween 80 and 15 µg mL^-1^ kanamycin, to grow overnight at 37°C with constant orbital shaking at 220 rpm. The starter culture was then used the next day to inoculate 5 L of Middlebrook 7H9 medium (USBiological) supplemented with 0.2% succinate, 0.05% Tween 80 and 15 µg mL^-1^ kanamycin, at an initial OD_600nm_ of 0.05, and left to grow overnight at 37°C with constant orbital shaking at 220 rpm. The next morning, AftD expression was induced by adding acetamide to a final concentration of 0.2%, once the culture reached an OD_600nm_ of 0.8 to 1.0, roughly 16 hr after inoculation. The culture was grown for 24 hr after induction at 37°C with constant orbital shaking at 220 rpm, then harvested at 4°C by centrifugation for 20 min at 4000 rpm (JA-10 rotor, Beckman Coulter). Supernatant was then discarded, and the cell pellet was washed once with PBS, centrifuged again at 4°C for 20 min at 4000 rpm, then frozen at −20°C.

The cells were thawed and resuspended in lysis buffer containing 20 mM HEPES pH 7.5, 200 mM μg/mL DNase I, 1 mM TCEP, 1 tablet/1.5 L buffer EDTA-free cOmplete protease inhibitor cocktail (Roche). For 5 L of culture, there was ∼10 g of wet cell pellet mass, which was resuspended in ∼60mL of lysis buffer. Cells were then lysed in a French Press, at 1500 bar, after 4 passages. To isolate the membrane fractions, the lysate was put through ultracentrifugation at 42,000g in Type 45 Ti Rotor (Beckman Coulter) at 4°C for 45 min. The supernatant was discarded, the pellet was resuspended in the lysis buffer up to a volume of 15 mL and homogenized using a hand-held glass homogenizer (Wheaton) and then stored at −20°C. The thawed membrane fraction was solubilized by adding DDM to a final concentration of 1% (w/v) detergent for 3 hours at 4°C with gentle rotation. Insoluble material was removed by ultracentrifugation at 42,000g in Type 45 Ti Rotor at 4°C for 45 min. The supernatant was added to a Falcon50 tube containing pre-equilibrated ANTI-FLAG® M2 Affinity Gel (Sigma) and incubated with gentle rotation overnight at 4°C. The gel was washed with 10 column volumes of wash buffer containing 20 mM HEPES pH 7.5, 200 mM NaCl, 0.05% DDM and incubated at 4°C with gentle rotation for 1h with elution buffer containing 20 mM HEPES pH 7.5, 200mM NaCl, 0.05% DDM, μg mL-1 FLAG peptide (Sigma), then finally eluted. The eluted fraction was then run on both SDS-PAGE and Native-PAGE gels and sent for subsequent mass spectrometry analysis.

### Mass spectrometry analysis and protein identification

Samples were subjected to trypsin digestion. Briefly, protein sample was reduced with 10 mM DTT (Sigma) for 40 min at 56 °C followed by alkylation with iodoacetamide (Sigma) 55 mM for 30 min in the dark. Excessive iodoacetamide was quenched by further incubation with DTT (10 mM for 10 min in the dark). The resulting sample was digested overnight with trypsin (Proteomics grade from Promega) at 37 °C (1:50 protein/trypsin ratio), dried and resuspended in 8 µL LCMS water 0.1% formic acid (Fisher Chemicals).

Nano-liquid chromatography-tandem mass spectrometry (nanoLC-MS/MS) analysis was performed on an ekspert™ NanoLC 425 cHiPLC® system coupled with a TripleTOF® 6600 with a NanoSpray® III source (Sciex). Peptides were separated through reversed-phase chromatography (RP-LC) in a trap-and-elute mode. Trapping was performed at 2 µl/min with 100% A (0.1% formic acid in water, Fisher Chemicals, Geel, Belgium), for 10 min, on a Nano cHiPLC Trap column (Sciex 200 µm x 0.5 mm, ChromXP C18-CL, 3 µm, 120 Å). Separation was performed at 300 nl/min, on a Nano cHiPLC column (Sciex 75 µm x 15 cm, ChromXP C18-CL, 3 µm, 120 Å). The gradient was as follows: 0-1 min, 5% B (0.1% formic acid in acetonitrile, Fisher Chemicals); 1-61 min, 5-30% B; 61-63 min, 30-80% B; 63-71 min, 80% B; 71-73 min, 80-5% B; 73-90 min, 5% B.

Peptides were sprayed into the MS through an uncoated fused-silica PicoTip™ emitter (360 µm O.D., 20 µm I.D., 10 ± 1.0 µm tip I.D., New Objective). The source parameters were set as follows: 15 GS1, 0 GS2, 30 CUR, 2.5 keV ISVF and 100 C IHT. An information dependent acquisition (IDA) method was set with a TOF-MS survey scan of 400-2000 m/z for 250 msec. The 50 most intense precursors were selected for subsequent fragmentation and the MS/MS were acquired in high sensitivity mode (150-1800 m/z for 40 msec each). The selection criteria for parent ions included a charge state between +2 and +5 and counts above a minimum threshold of 125 counts per second. Ions were excluded from further MSMS analysis for 12 s. Fragmentation was performed using rolling collision energy with a collision energy spread of 5.

The obtained spectra were processed and analyzed using ProteinPilot™ software, with the Paragon search engine (version 5.0, Sciex). The following search parameters were set: search against *M. smegmatis* and *E. coli* from Uniprot/SwissProt database, and protein sequence of AftD from *M. abscessus*; Iodoacetamide, as Cys alkylation; Tryspsin, as digestion; TripleTOF 6600, as the Instrument; ID focus as biological modifications and Amino acid substitutions; search effort as thorough; and a FDR analysis. Only the proteins with Unused Protein Score above 1.3 and 95% confidence were considered.

## REFERENCES

Abrahams, K.A., and Besra, G.S. (2016). Mycobacterial cell wall biosynthesis: a multifaceted antibiotic target. Parasitology, 1–18.

Adams, P.D., Afonine, P.V., Bunkóczi, G., Chen, V.B., Davis, I.W., Echols, N., Headd, J.J., Hung, L.-W., Kapral, G.J., and Grosse-Kunstleve, R.W. (2010). PHENIX: a comprehensive Python-based system for macromolecular structure solution. Acta Crystallographica Section D: Biological Crystallography 66, 213–221.

Afonine, P.V., Poon, B.K., Read, R.J., Sobolev, O.V., Terwilliger, T.C., Urzhumtsev, A., and Adams, P.D. (2018). Real-space refinement in PHENIX for cryo-EM and crystallography. Acta Crystallographica Section D: Structural Biology 74.

Alderwick, L., Birch, H., Mishra, A., Eggeling, L., and Besra, G. (2007). Structure, function and biosynthesis of the Mycobacterium tuberculosis cell wall: arabinogalactan and lipoarabinomannan assembly with a view to discovering new drug targets (Portland Press Limited).

Alderwick, L.J., Birch, H.L., Krumbach, K., Bott, M., Eggeling, L., and Besra, G.S. (2018). AftD functions as an *α* 1→5 arabinofuranosyltransferase involved in the biosynthesis of the mycobacterial cell wall core. The Cell Surface 1, 2–14.

Alderwick, L.J., Harrison, J., Lloyd, G.S., and Birch, H.L. (2015). The Mycobacterial cell wall— Peptidoglycan and arabinogalactan. Cold Spring Harbor perspectives in medicine, a021113.

Alderwick, L.J., Seidel, M., Sahm, H., Besra, G.S., and Eggeling, L. (2006). Identification of a novel arabinofuranosyltransferase (AftA) involved in cell wall arabinan biosynthesis in Mycobacterium tuberculosis. Journal of Biological Chemistry 281, 15653–15661.

Alsteens, D., Verbelen, C., Dague, E., Raze, D., Baulard, A.R., and Dufrêne, Y.F. (2008). Organization of the mycobacterial cell wall: a nanoscale view. Pflügers Archiv-European Journal of Physiology 456, 117–125.

Altschul, S.F., and Koonin, E.V. (1998). Iterated profile searches with PSI-BLAST—a tool for discovery in protein databases. Trends in biochemical sciences 23, 444–447.

Angala, S.k., McNeil, M.R., Zou, L., Liav, A., Zhang, J., Lowary, T.L., and Jackson, M. (2016). Identification of a novel mycobacterial arabinosyltransferase activity which adds an arabinosyl α-D-mannosyl residues. ACS chemical biology 11, 1518–1524.

Bai, L., Kovach, A., You, Q., Kenny, A., and Li, H. (2019). Structure of the eukaryotic protein O-mannosyltransferase Pmt1− Pmt2 complex. Nature structural & molecular biology 26, 704–711.

Bai, L., Wang, T., Zhao, G., Kovach, A., and Li, H. (2018). The atomic structure of a eukaryotic oligosaccharyltransferase complex. Nature 555, 328.

Bai, X.-c., Rajendra, E., Yang, G., Shi, Y., and Scheres, S.H. (2015). Sampling the conformational space of the catalytic subunit of human γ

Barad, B.A., Echols, N., Wang, R.Y.-R., Cheng, Y., DiMaio, F., Adams, P.D., and Fraser, J.S. (2015). EMRinger: side chain–directed model and map validation for 3D cryo-electron microscopy. Nature methods 12, 943.

Barry, C.E., Crick, D.C., and McNeil, M.R. (2007). Targeting the formation of the cell wall core of M. tuberculosis. Infectious Disorders-Drug Targets (Formerly Current Drug Targets-Infectious Disorders) 7, 182–202.

Barry III, C.E. (2001). Interpreting cell wall ‘virulence factors’ of Mycobacterium tuberculosis. Trends in microbiology 9, 237–241.

Battesti, A., and Bouveret, E. (2009). Bacteria possessing two RelA/SpoT-like proteins have evolved a specific stringent response involving the acyl carrier protein-SpoT interaction. Journal of bacteriology 191, 616–624.

Bayburt, T.H., and Sligar, S.G. (2010). Membrane protein assembly into Nanodiscs. FEBS letters 584, 1721–1727.

Belanger, A.E., Besra, G.S., Ford, M.E., Mikusová, K., Belisle, J.T., Brennan, P.J., and Inamine, J.M. (1996). The embAB genes of Mycobacterium avium encode an arabinosyl transferase involved in cell wall arabinan biosynthesis that is the target for the antimycobacterial drug ethambutol. Proceedings of the National Academy of Sciences 93, 11919–11924.

Birch, H.L., Alderwick, L.J., Bhatt, A., Rittmann, D., Krumbach, K., Singh, A., Bai, Y., Lowary, T.L., Eggeling, L., and Besra, G.S. (2008). Biosynthesis of mycobacterial arabinogalactan: identification of a novel *α* (1→3) arabinofuranosyltransferase. Molecular microbiology 69, 1191–1206.

Boraston, A.B., Bolam, D.N., Gilbert, H.J., and Davies, G.J. (2004). Carbohydrate-binding modules: fine-tuning polysaccharide recognition. Biochemical journal 382, 769–781.

Boraston, A.B., Kwan, E., Chiu, P., Warren, R.A.J., and Kilburn, D.G. (2003). Recognition and hydrolysis of noncrystalline cellulose. Journal of biological chemistry 278, 6120–6127.

Bruni, R., and Kloss, B. (2013). High - Throughput Cloning and Expression of Integral Membrane Proteins in Escherichia coli. Current protocols in protein science 74, 29.26. 21–29.26. 34.

Burnley, T., Palmer, C.M., and Winn, M. (2017). Recent developments in the CCP - EM software suite. Acta Crystallographica Section D 73, 469–477.

Catalão, M.J., Gil, F., Moniz-Pereira, J., and Pimentel, M. (2010). The mycobacteriophage Ms6 encodes a chaperone like protein involved in the endolysin delivery to the peptidoglycan. Molecular microbiology 77, 672–686.

Chen, V.B., Arendall, W.B., Headd, J.J., Keedy, D.A., Immormino, R.M., Kapral, G.J., Murray, L.W., Richardson, J.S., and Richardson, D.C. (2010). MolProbity: all-atom structure validation for macromolecular crystallography. Acta Crystallographica Section D: Biological Crystallography 66, 12–21.

Cheng, A., Eng, E.T., Alink, L., Rice, W.J., Jordan, K.D., Kim, L.Y., Potter, C.S., and Carragher, B. (2018). High resolution single particle cryo-electron microscopy using beam-image shift. Journal of Structural Biology.

Cohen, K.A., El-Hay, T., Wyres, K.L., Weissbrod, O., Munsamy, V., Yanover, C., Aharonov, R., Shaham, O., Conway, T.C., and Goldschmidt, Y. (2016). Paradoxical hypersusceptibility of drug resistant mycobacteriumtuberculosis to β-lactam antibiotics. EBioMedicine 9, 170–179.

Cowtan, K. (2006). The Buccaneer software for automated model building. 1. Tracing protein chains. Acta Crystallographica Section D: Biological Crystallography 62, 1002–1011.

Cronan, J.E. (2014). The chain-flipping mechanism of ACP (acyl carrier protein)-dependent enzymes appears universal. Biochemical Journal 460, 157–163.

Dandey, V.P., Wei, H., Zhang, Z., Tan, Y.Z., Acharya, P., Eng, E.T., Rice, W.J., Kahn, P.A., Potter, C.S., and Carragher, B. (2018). Spotiton: New features and applications. Journal of Structural Biology 202, 161–169.

DeJesus, M.A., Gerrick, E.R., Xu, W., Park, S.W., Long, J.E., Boutte, C.C., Rubin, E.J., Schnappinger, D., Ehrt, S., and Fortune, S.M. (2017). Comprehensive essentiality analysis of the Mycobacterium tuberculosis genome via saturating transposon mutagenesis. MBio 8, e02133–02116.

Edgar, R.C. (2004). MUSCLE: multiple sequence alignment with high accuracy and high throughput. Nucleic acids research 32, 1792–1797.

Emsley, P., and Cowtan, K. (2004). Coot: model-building tools for molecular graphics. Acta Crystallographica Section D: Biological Crystallography 60, 2126–2132.

Favrot, L., and Ronning, D.R. (2012). Targeting the mycobacterial envelope for tuberculosis drug development. Expert review of anti-infective therapy 10, 1023–1036.

Goude, R., Amin, A., Chatterjee, D., and Parish, T. (2009). The arabinosyltransferase EmbC is inhibited by ethambutol in Mycobacterium tuberculosis. Antimicrobial agents and chemotherapy 53, 4138–4146.

Grant, T., Rohou, A., and Grigorieff, N. (2018). cisTEM, user-friendly software for single-particle image processing. Elife 7, e35383.

Grzegorzewicz, A.E., De Sousa-d’Auria, C., McNeil, M.R., Huc-Claustre, E., Jones, V., Petit, C., Angala, S.K., Zemanová, J., Wang, Q., and Belardinelli, J.M. (2016). Assembling of the Mycobacterium tuberculosis cell wall core. Journal of Biological Chemistry, jbc. M116. 739227.

Hoffmann, C., Leis, A., Niederweis, M., Plitzko, J.M., and Engelhardt, H. (2008). Disclosure of the mycobacterial outer membrane: cryo-electron tomography and vitreous sections reveal the lipid bilayer structure. Proceedings of the National Academy of Sciences 105, 3963–3967.

Holm, L., and Laakso, L.M. (2016). Dali server update. Nucleic acids research 44, W351–W355.

Islam, M.M., Hameed, H.A., Mugweru, J., Chhotaray, C., Wang, C., Tan, Y., Liu, J., Li, X., Tan, S., and Ojima, I. (2017). Drug resistance mechanisms and novel drug targets for tuberculosis therapy. Journal of genetics and genomics 44, 21–37.

Jain, T., Sheehan, P., Crum, J., Carragher, B., and Potter, C.S. (2012). Spotiton: a prototype for an integrated inkjet dispense and vitrification system for cryo-TEM. Journal of structural biology 179, 68–75.

Jankute, M., Cox, J.A., Harrison, J., and Besra, G.S. (2015). Assembly of the mycobacterial cell wall. Annual review of microbiology 69, 405–423.

Jena, L., Nayak, T., Deshmukh, S., Wankhade, G., Waghmare, P., and Harinath, C. (2016). Isoniazid with multiple mode of action on various mycobacterial enzymes resulting in drug resistance. J Infect Dis Ther 4, 2332–0877.1000297.

Joe, M., and Lowary, T.L. (2006). Synthesis of 2-deoxy-2-fluoro analogs of polyprenyl β-D-arabinofuranosyl phosphates. Carbohydrate research 341, 2723–2730.

Kapust, R.B., Tözsér, J., Copeland, T.D., and Waugh, D.S. (2002). The P1′ specificity of tobacco etch virus protease. Biochemical and biophysical research communications 294, 949–955.

Kim, J.-H., O’Brien, K.M., Sharma, R., Boshoff, H.I., Rehren, G., Chakraborty, S., Wallach, J.B., Monteleone, M., Wilson, D.J., and Aldrich, C.C. (2013). A genetic strategy to identify targets for the development of drugs that prevent bacterial persistence. Proceedings of the National Academy of Sciences 110, 19095–19100.

Kimanius, D., Forsberg, B.O., Scheres, S.H., and Lindahl, E. (2016). Accelerated cryo-EM structure determination with parallelisation using GPUs in RELION-2. Elife 5, e18722.

Kordulakova, J., Gilleron, M., Mikusová, K.n., Puzo, G., Brennan, P.J., Gicquel, B., and Jackson, M. (2002). Definition of the first mannosylation step in phosphatidylinositol mannoside synthesis PimA is essential for growth of mycobacteria. Journal of Biological Chemistry 277, 31335–31344.

Kreamer, N.N., Chopra, R., Caughlan, R.E., Fabbro, D., Fang, E., Gee, P., Hunt, I., Li, M., Leon, B.C., and Muller, L. (2018). Acylated-acyl carrier protein stabilizes the Pseudomonas aeruginosa WaaP lipopolysaccharide heptose kinase. Scientific reports 8, 14124.

Kremer, L., Nampoothiri, K.M., Lesjean, S., Dover, L.G., Graham, S., Betts, J., Brennan, P.J., Minnikin, D.E., Locht, C., and Besra, G.S. (2001). Biochemical Characterization of Acyl Carrier Protein (AcpM) and Malonyl-CoA: AcpM Transacylase (mtFabD), Two Major Components ofMycobacterium tuberculosis Fatty Acid Synthase II. Journal of Biological Chemistry 276, 27967–27974.

Krogh, A., Larsson, B., Von Heijne, G., and Sonnhammer, E.L. (2001). Predicting transmembrane protein topology with a hidden Markov model: application to complete genomes. Journal of molecular biology 305, 567–580.

Kucukelbir, A., Sigworth, F.J., and Tagare, H.D. (2013). Quantifying the local resolution of cryo-EM density maps. Nature methods 11, 63.

Lairson, L., Henrissat, B., Davies, G., and Withers, S. (2008). Glycosyltransferases: structures, functions, and mechanisms. Annual review of biochemistry 77.

Lander, G.C., Stagg, S.M., Voss, N.R., Cheng, A., Fellmann, D., Pulokas, J., Yoshioka, C., Irving, C., Mulder, A., Lau, P.W., et al. (2009). Appion: an integrated, database-driven pipeline to facilitate EM image processing. J Struct Biol 166, 95–102.

Layre, E., Al-Mubarak, R., Belisle, J.T., and Moody, D.B. (2014). Mycobacterial lipidomics. In Molecular Genetics of Mycobacteria, Second Edition (American Society of Microbiology), pp. 341–360.

Lea-Smith, D.J., Martin, K.L., Pyke, J.S., Tull, D., McConville, M.J., Coppel, R.L., and Crellin, P.K. (2008). Analysis of a new mannosyltransferase required for the synthesis of phosphatidylinositol mannosides and lipoarbinomannan reveals two lipomannan pools in corynebacterineae. Journal of Biological Chemistry 283, 6773–6782.

Li, W., and Godzik, A. (2006). Cd-hit: a fast program for clustering and comparing large sets of protein or nucleotide sequences. Bioinformatics 22, 1658–1659.

Liu, J., and Mushegian, A. (2003). Three monophyletic superfamilies account for the majority of the known glycosyltransferases. Protein Science 12, 1418–1431.

Lizak, C., Gerber, S., Numao, S., Aebi, M., and Locher, K.P. (2011). X-ray structure of a bacterial oligosaccharyltransferase. Nature 474, 350.

Love, J., Mancia, F., Shapiro, L., Punta, M., Rost, B., Girvin, M., Wang, D.-N., Zhou, M., Hunt, J.F., and Szyperski, T. (2010). The New York Consortium on Membrane Protein Structure (NYCOMPS): a high-throughput platform for structural genomics of integral membrane proteins. Journal of structural and functional genomics 11, 191–199.

Ludtke, S.J., Baldwin, P.R., and Chiu, W. (1999). EMAN: semiautomated software for high-resolution single-particle reconstructions. Journal of structural biology 128, 82–97.

Ludwig, W., Euzéby, J., Schumann, P., Busse, H.J., Trujillo, M.E., Kämpfer, P., and Whitman, W.B. (2015). Road map of the phylum Actinobacteria. Bergey’s Manual of Systematics of Archaea and Bacteria, 1–37.

Majerus, P.W., Alberts, A.W., and Vagelos, P.R. (1965). Acyl carrier protein, IV. The identification of 4′-phosphopantetheine as the prosthetic group of the acyl carrier protein. Proceedings of the National Academy of Sciences 53, 410–417.

Mancia, F., and Love, J. (2010). High-throughput expression and purification of membrane proteins. Journal of structural biology 172, 85–93.

Matsumoto, S., Shimada, A., Nyirenda, J., Igura, M., Kawano, Y., and Kohda, D. (2013). Crystal structures of an archaeal oligosaccharyltransferase provide insights into the catalytic cycle of N-linked protein glycosylation. Proceedings of the National Academy of Sciences 110, 17868–17873.

Matsumoto, S., Taguchi, Y., Shimada, A., Igura, M., and Kohda, D. (2017). Tethering an N-glycosylation sequon-containing peptide creates a catalytically competent oligosaccharyltransferase complex. Biochemistry 56, 602–611.

Morita, Y.S., Sena, C.B., Waller, R.F., Kurokawa, K., Sernee, M.F., Nakatani, F., Haites, R.E., Billman-Jacobe, H., McConville, M.J., and Maeda, Y. (2006). PimE is a polyprenol-phosphate-mannose-dependent mannosyltransferase that transfers the fifth mannose of phosphatidylinositol mannoside in mycobacteria. Journal of Biological Chemistry.

Napiórkowska, M., Boilevin, J., Darbre, T., Reymond, J.-L., and Locher, K.P. (2018). Structure of bacterial oligosaccharyltransferase PglB bound to a reactive LLO and an inhibitory peptide. Scientific reports 8, 16297.

Napiórkowska, M., Boilevin, J., Sovdat, T., Darbre, T., Reymond, J.-L., Aebi, M., and Locher, K.P. (2017). Molecular basis of lipid-linked oligosaccharide recognition and processing by bacterial oligosaccharyltransferase. Nature structural & molecular biology 24, 1100.

Nguyen, C., Haushalter, R.W., Lee, D.J., Markwick, P.R., Bruegger, J., Caldara-Festin, G., Finzel, K., Jackson, D.R., Ishikawa, F., and O’dowd, B. (2014). Trapping the dynamic acyl carrier protein in fatty acid biosynthesis. Nature 505, 427.

Nikaido, H., and Jarlier, V. (1991). Permeability of the mycobacterial cell wall. Research in microbiology 142, 437–443.

Ofer, N., Wishkautzan, M., Meijler, M., Wang, Y., Speer, A., Niederweis, M., and Gur, E. (2012). Ectoine biosynthesis in *Mycobacterium smegmatis*. Appl Environ Microbiol 78, 7483–7486.

Petrou, V.I., Herrera, C.M., Schultz, K.M., Clarke, O.B., Vendome, J., Tomasek, D., Banerjee, S., Rajashankar, K.R., Dufrisne, M.B., and Kloss, B. (2016). Structures of aminoarabinose transferase ArnT suggest a molecular basis for lipid A glycosylation. Science 351, 608–612.

Pettersen, E.F., Goddard, T.D., Huang, C.C., Couch, G.S., Greenblatt, D.M., Meng, E.C., and Ferrin, T.E. (2004). UCSF Chimera—a visualization system for exploratory research and analysis. Journal of computational chemistry 25, 1605–1612.

Portevin, D., de Sousa-D’Auria, C., Houssin, C., Grimaldi, C., Chami, M., Daffé, M., and Guilhot, C.(2004). A polyketide synthase catalyzes the last condensation step of mycolic acid biosynthesis in mycobacteria and related organisms. Proceedings of the National Academy of Sciences 101, 314–319.

Punjani, A., Rubinstein, J.L., Fleet, D.J., and Brubaker, M.A. (2017). cryoSPARC: algorithms for rapid unsupervised cryo-EM structure determination. Nat Methods 14, 290–296.

Razinkov, I., Dandey, V.P., Wei, H., Zhang, Z., Melnekoff, D., Rice, W.J., Wigge, C., Potter, C.S., and Carragher, B. (2016). A new method for vitrifying samples for cryoEM. Journal of structural biology 195, 190–198.

Rohou, A., and Grigorieff, N. (2015). CTFFIND4: Fast and accurate defocus estimation from electron micrographs. J Struct Biol 192, 216–221.

Rosenthal, P.B., and Henderson, R. (2003). Optimal determination of particle orientation, absolute hand, and contrast loss in single-particle electron cryomicroscopy. J Mol Biol 333, 721–745.

Schaeffer, M.L., Agnihotri, G., Kallender, H., Brennan, P.J., and Lonsdale, J.T. (2001). Expression, purification, and characterization of the Mycobacterium tuberculosis acyl carrier protein, AcpM. Biochimica et Biophysica Acta (BBA)-Molecular and Cell Biology of Lipids 1532, 67–78.

Scheres, S.H. (2012). RELION: implementation of a Bayesian approach to cryo-EM structure determination. J Struct Biol 180, 519–530.

Seidel, M., Alderwick, L.J., Birch, H.L., Sahm, H., Eggeling, L., and Besra, G.S. (2007). Identification of a novel arabinofuranosyltransferase AftB involved in a terminal step of cell wall arabinan biosynthesis in Corynebacterianeae, such as Corynebacterium glutamicum and Mycobacterium tuberculosis. Journal of Biological Chemistry.

Sharma, C.B., Lehle, L., and Tanner, W. (1981). N - Glycosylation of Yeast Proteins: Characterization of the Solubilized Oligosaccharyl Transferase. European journal of biochemistry 116, 101–108.

Shekar, S., Yeo, Z.X., Wong, J.C., Chan, M.K., Ong, D.C., Tongyoo, P., Wong, S.-Y., and Lee, A.S. (2014). Detecting novel genetic variants associated with isoniazid-resistant Mycobacterium tuberculosis. PloS one 9, e102383.

Škovierová, H., Larrouy-Maumus, G., Zhang, J., Kaur, D., Barilone, N., Korduláková, J., Gilleron, M., Guadagnini, S., Belanová, M., and Prevost, M.-C. (2009). AftD, a novel essential arabinofuranosyltransferase from mycobacteria. Glycobiology 19, 1235–1247.

Stewart, G.R., Wernisch, L., Stabler, R., Mangan, J.A., Hinds, J., Laing, K.G., Butcher, P.D., and Young, D.B. (2002). The heat shock response of Mycobacterium tuberculosis: linking gene expression, immunology and pathogenesis. International Journal of Genomics 3, 348–351.

Suloway, C., Pulokas, J., Fellmann, D., Cheng, A., Guerra, F., Quispe, J., Stagg, S., Potter, C.S., and Carragher, B. (2005). Automated molecular microscopy: the new Leginon system. Journal of structural biology 151, 41–60.

Takayama, K., and Kilburn, J.O. (1989). Inhibition of synthesis of arabinogalactan by ethambutol in Mycobacterium smegmatis. Antimicrobial agents and chemotherapy 33, 1493–1499.

Takayama, K., Wang, C., and Besra, G.S. (2005). Pathway to synthesis and processing of mycolic acids in Mycobacterium tuberculosis. Clinical microbiology reviews 18, 81–101.

Tallorin, L., Finzel, K., Nguyen, Q.G., Beld, J., La Clair, J.J., and Burkart, M.D. (2016). Trapping of the enoyl-acyl carrier protein reductase–acyl carrier protein interaction. Journal of the American Chemical Society 138, 3962–3965.

Tan, Y.Z., Baldwin, P.R., Davis, J.H., Williamson, J.R., Potter, C.S., Carragher, B., and Lyumkis, (2017). Addressing preferred specimen orientation in single-particle cryo-EM through tilting. Nature methods 14, 793.

Telenti, A., Philipp, W.J., Sreevatsan, S., Bernasconi, C., Stockbauer, K.E., Wieles, B., Musser, J.M., and Jacobs Jr, W.R. (1997). The emb operon, a gene cluster of Mycobacterium tuberculosis involved in resistance to ethambutol. Nature medicine 3, 567.

Therisod, H., and Kennedy, E.P. (1987). The function of acyl carrier protein in the synthesis of membrane-derived oligosaccharides does not require its phosphopantetheine prosthetic group. Proceedings of the National Academy of Sciences 84, 8235–8238.

Voss, N.R., and Gerstein, M. (2010). 3V: cavity, channel and cleft volume calculator and extractor. Nucleic Acids Res 38, W555–562.

Wallace, A.C., Laskowski, R.A., and Thornton, J.M. (1995). LIGPLOT: a program to generate schematic diagrams of protein-ligand interactions. Protein engineering, design and selection 8, 127–134.

Wei, H., Dandey, V.P., Zhang, Z., Raczkowski, A., Rice, W.J., Carragher, B., and Potter, C.S. (2018). Optimizing “self-wicking” nanowire grids. Journal of structural biology 202, 170–174.

WHO (2018). Global tuberculosis report 2018.

Wild, R., Kowal, J., Eyring, J., Ngwa, E.M., Aebi, M., and Locher, K.P. (2018). Structure of the yeast oligosaccharyltransferase complex gives insight into eukaryotic N-glycosylation. Science 359, 545–550.

Wolucka, B.A., McNeil, M.R., de Hoffmann, E., Chojnacki, T., and Brennan, P.J. (1994). Recognition of the lipid intermediate for arabinogalactan/arabinomannan biosynthesis and its relation to the mode of action of ethambutol on mycobacteria. Journal of Biological Chemistry 269, 23328–23335.

Wong, H.C., Liu, G., Zhang, Y.-M., Rock, C.O., and Zheng, J. (2002). The solution structure of acyl carrier protein from Mycobacterium tuberculosis. Journal of Biological Chemistry 277, 15874–15880.

Zhang, K. (2016). Gctf: Real-time CTF determination and correction. J Struct Biol 193, 1–12.

Zheng, R.B., Jégouzo, S.A., Joe, M., Bai, Y., Tran, H.-A., Shen, K., Saupe, J.r., Xia, L., Ahmed, M.F., and Liu, Y.-H. (2017a). Insights into interactions of mycobacteria with the host innate immune system from a novel array of synthetic mycobacterial glycans. ACS chemical biology 12, 2990–3002.

Zheng, S.Q., Palovcak, E., Armache, J.P., Verba, K.A., Cheng, Y., and Agard, D.A. (2017b). MotionCor2: anisotropic correction of beam-induced motion for improved cryo-electron microscopy. Nat Methods 14, 331–332.

Zimhony, O., Schwarz, A., Raitses-Gurevich, M., Peleg, Y., Dym, O., Albeck, S., Burstein, Y., and Shakked, Z. (2015). AcpM, the meromycolate extension acyl carrier protein of Mycobacterium tuberculosis, is activated by the 4′-phosphopantetheinyl transferase PptT, a potential target of the multistep mycolic acid biosynthes is. Biochemistry 54, 2360–2371.

